# RNADiffFold: Generative RNA Secondary Structure Prediction using Discrete Diffusion Models

**DOI:** 10.1101/2024.05.28.596177

**Authors:** Zhen Wang, Yizhen Feng, Qingwen Tian, Ziqi Liu, Pengju Yan, Xiaolin Li

**Affiliations:** Hangzhou Institute of Medicine, Chinese Academy of Sciences, Hangzhou, 310018, Zhejiang, China; College of Information Engineering, Zhejiang University of Technology, Hangzhou, 310014, Zhejiang, China; College of Computer Science and Technology, Zhejiang University of Technology, Hangzhou, 310014, Zhejiang, China; Hangzhou Institute for Advanced Study, University of Chinese Academy of Sciences, Hangzhou, 310024, Zhejiang, China

**Keywords:** RNA secondary structure prediction, deep learning, discrete diffusion model

## Abstract

RNA molecules are essential macromolecules that perform diverse biological functions in living beings. Precise prediction of RNA secondary structures is instrumental in deciphering their complex three-dimensional architecture and functionality. Traditional methodologies for RNA structure prediction, including energy-based and learning-based approaches, often depict RNA secondary structures from a static perspective and rely on stringent a priori constraints. Inspired by the success of diffusion models, in this work, we introduce RNADiffFold, an innovative generative prediction approach of RNA secondary structures based on multinomial diffusion. We reconceptualize the prediction of contact maps as akin to pixel-wise segmentation and accordingly train a denoising model to refine the contact maps starting from a noise-infused state progressively. We also devise a potent conditioning mechanism that harnesses features extracted from RNA sequences to steer the model toward generating an accurate secondary structure. These features encompass one-hot encoded sequences, probabilistic maps generated from a pre-trained scoring network, and embeddings and attention maps derived from RNA-FM. Experimental results on both within- and cross-family datasets demonstrate RNADiffFold’s competitive performance compared with current state-of-the-art methods. Additionally, RNADiffFold has shown a notable proficiency in capturing the dynamic aspects of RNA structures, a claim corroborated by its performance on datasets comprising multiple conformations.

## Introduction

Ribonucleic acid (RNA) is a vital biomolecule with diverse roles beyond the transfer of genetic information from DNA to proteins. It is involved in catalysis, regulation, and protein synthesis, primarily through its non-coding regions [1]. For instance, microRNAs (miRNAs) regulate gene expression post-transcriptionally, with their dysregulation linked to various diseases [2]. Long non-coding RNAs (lncRNAs) are crucial in cellular processes like chromatin modification and transcriptional regulation [3]. Small nuclear RNAs (snRNAs) form spliceosome complexes, influencing splicing patterns and mRNA abundance [4]. Thus, delving into RNA functionality is essential for unraveling its biological mechanisms.

In cells, RNA typically exists as a single-stranded molecule and folds into specific structures through base pairing via hydrogen bonds, thereby interacting with other biomolecules and exerting functions. The secondary structure of RNA is a two-dimensional topological structure formed by base pairing through hydrogen bonds [5]. Based on this, the tertiary structure further folds into a three-dimensional spatial conformation. Although RNA primarily exerts its biological functions through its tertiary structure, this binding process often relies on motifs such as stems and loops in the secondary structure. However, due to its single-stranded nature, RNA’s tertiary structure is susceptible to environmental influences and exhibits poor stability, making it challenging to obtain directly. Despite significant progress in resolving high-resolution RNA structures through existing experimental methods such as X-ray crystallography, nuclear magnetic resonance (NMR) [6], and cryo-electron microscopy [7], challenges persist in obtaining a sufficient quantity and quality of RNA tertiary structures due to factors such as high experimental costs and resolution limitations. Therefore, understanding RNA’s secondary structure is key to deciphering its biological mechanisms and progressing to tertiary structure resolution [8].

Over the past decades, researchers have developed various methods for RNA secondary structure prediction, combining experimental and computational approaches. These methods fall into three main categories: energy-based, covariation-based, and deep learning-based [9]. Energy-based methods primarily utilize experimentally determined parameters to calculate the free energy of RNA structures and identify the most stable secondary structures through dynamic programming [10, 11, 12, 13]. While widely adopted and capable of providing relatively accurate predictions, these methods are limited in that they only consider nested base pairings and struggle with more complex structures such as pseudoknots. Moreover, as the length of RNA increases, the computational complexity of these methods significantly escalates. Covariation-based methods infer secondary structures by considering the co-evolutionary relationships between RNA sequences and structures [14, 15, 16]. These methods can offer highly accurate predictions under certain conditions but may face challenges when dealing with limited information from homologous sequence data. With the accumulation of RNA secondary structure data and the rapid development of deep learning technologies, deep learning-based RNA secondary structure prediction methods have gained traction. These methods employ neural networks, such as Bi-LSTM, Transformer, and Unet to calculate base pairing probabilities to capture long-range interactions [17, 18, 19]. Some approaches integrate thermodynamic knowledge [20], adopt transfer learning strategies [21], or incorporate evolutionary and mutational coupling information [22] to optimize prediction results and alleviate prediction biases. However, existing deep learning-based methods still exhibit shortcomings in model generalization performance, especially when modeling unknown RNA families. This limitation arises because their model parameters often derive from a limited pool of known structures, thereby restricting their adaptability to new data. To overcome limitations in training data quantity and distribution and enhance model generalization capabilities, new computational methods are necessary to achieve more accurate and comprehensive outcomes in RNA secondary structure prediction.

In recent years, diffusion models have demonstrated outstanding performance in various prediction tasks [23, 24, 25]. Inspired by this, we introduce **RNADiffFold**, a novel framework for RNA secondary structure prediction using discrete diffusion models. The framework aims to predict a deterministic RNA secondary structure in a generative manner. RNADiffFold first represents RNA secondary structures as binary contact maps. The contact map has a size of *L×L* (where L is the length of the RNA sequence), and each point in the map can be classified into two categories: “1” denotes pairing, and “0” denotes non-pairing. This approach simplifies the complex RNA secondary structure prediction task into a pixel-level image segmentation task.

RNADiffFold comprises two main components: the diffusion model and conditional control. The diffusion model component is based on discrete data space multinomial diffusion [26]. As illustrated in Figure 1, during the forward diffusion process, the true contact map *x*_0_ is gradually degraded by injecting noise following a uniform categorical distribution. When reaching time step *T*, *x_T_* transitions into a completely random noise state. In the inverse diffusion process, we employ UNet [27] as the learning network and add conditional control to gradually denoise and restore the original contact map. The conditional control component encompasses the sequence information of RNA, including features such as one-hot encoding of the sequence, probability maps from the Ufold scoring network [19], and high-dimensional embeddings and attention maps from RNA-FM [28], with dimensionality reduction through different MLPs. At each time step of the inverse diffusion process, all these sequence features are fused with the intermediate state *x_t_*. This design enables RNADiffFold to leverage the powerful capabilities of the diffusion model to predict RNA secondary structures while integrating various sequence features to enhance the accuracy and stability of predictions.

**Fig. 1.**
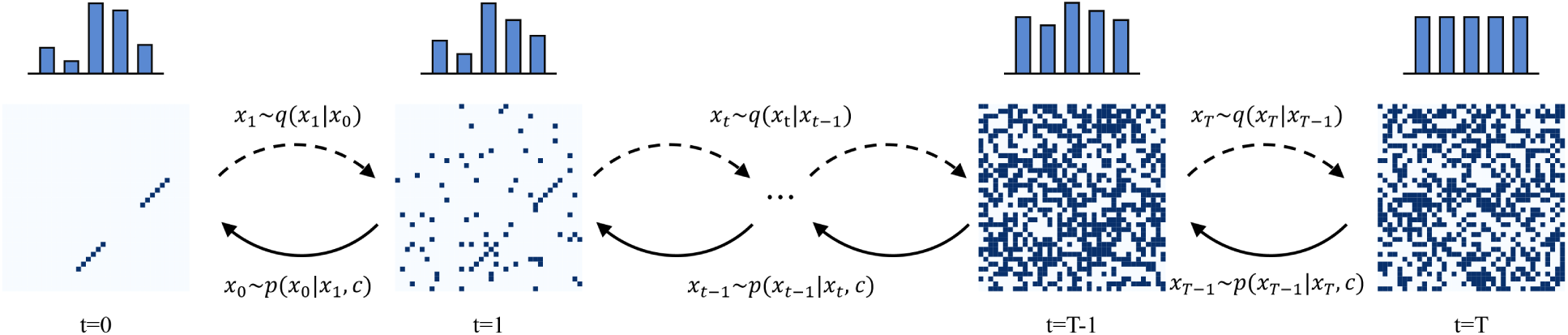
Overview of RNA secondary structure prediction with multinomial diffusion. During the diffusion process *q*(*x_t_|x_t−_*_1_), discrete noise is gradually introduced to corrupt the contact map from left to right. In the denoise process *p_θ_* (*x_t−_*_1_*|x_t_, c*), a generative model learns to denoise the corrupted map from right to left.

Building upon previous work [19, 21, 22], we evaluate the predictive performance of RNADiffFold on both within-family and cross-family RNA datasets. Experimental results demonstrate that even with simple one-hot encoding as the conditional input, RNADiffFold exhibits comparable performance to existing methods on within-family datasets, while also demonstrating reasonable accuracy in predicting the secondary structures of RNA sequences from unknown families. When additional features generated from RNA-FM are incorporated, RNADiffFold shows significant improvements in performance on both within-family and cross-family datasets, surpassing existing methods and highlighting its effectiveness and generalization. Furthermore, RNADiffFold not only predicts static RNA secondary structures but also captures dynamic multi-conformational features to some extent by learning the distribution of secondary structure conformations.

## Materials and methods

### Preliminaries

#### Diffusion models

Diffusion models [29, 30, 31] are a type of probabilistic generative models characterized by two Markov chains in the diffusion process: a forward chain that deconstructs data into noise, and a reverse chain that reconstructs data from noise. Specifically, in Diffusion Probabilistic Models (DDPMs), given a data distribution *x*_0_ *∼ q*(*x*_0_), the forward diffusion process *q* produces a sequence of latent states from *x*_1_ to *x_T_* by adding noise at the timestep *t* with variance schedule *β_t_ ∈*(0, 1). The transition kernel is defined as follows:

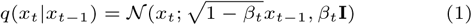

The reverse diffusion process *p* is parameterized by a prior distribution *x_T_ ∼ N* (0, **I**) and a learnable transition kernel *p_θ_*(*x_t−_*_1_*|x_t_*) defined as follows:

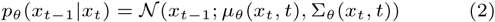

where *θ* denotes the model parameters, *µ_θ_* and Σ*_θ_* represent the mean and variance of the distribution at time *t*. The training objective is to learn the parameters *θ* so that the reserve trajectory *p_θ_* closely approximates the forward trajectory *q*. It is achieved by optimizing a variational upper bound on the negative log-likelihood:

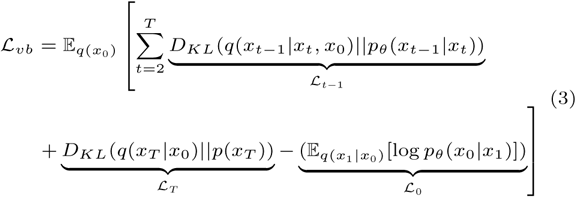

Due to the incorporation of Gaussian noise as a prior, most diffusion models operate effectively in continuous state spaces; however, they may not efficiently handle discrete data. In addressing this issue, some methods [32, 26, 33] are proposed to generate high-dimension discrete data. For example, D3PM [32] considers a transition matrix with an absorbing state or using discretized, truncated Gaussian distribution. VQ-diffusion [33] accommodates discrete data with a lazy random walk or a random masking operation.

#### RNA Foundation Model

RNA-FM (RNA Foundation Model) [28] is an RNA foundational model based on the BERT language model architecture, built upon 12 bidirectional encoder blocks based on Transformers. The model consists of two stages: pre-training and fine-tuning. In the pre-training stage, RNA-FM is trained in a self-supervised manner on a large amount of unlabeled RNA sequence data, allowing it to capture latent structural and functional information and extract meaningful RNA representations. Its pre-training strategy is similar to BERT, where 15% of the base tokens representing nucleotides are randomly masked, and the model is trained to reconstruct the masked tokens from the remaining sequence. Upon completion of training, RNA-FM can generate a 640 *× L* embedding matrix for each RNA sequence of length *L*. These embedding matrices provide rich feature representations for downstream tasks. In the task-specific fine-tuning stage, the pre-trained RNA-FM model can generate sequence embeddings tailored to the requirements of downstream modules, which can be directly used for various RNA-related machine-learning tasks.

It is worth noting that a recent study [34] further validated the importance of the multi-head attention mechanism outputs in RNA-FM for capturing structural information. These attention maps not only reveal the strength of associations between different positions in RNA sequences but also provide new perspectives for understanding RNA secondary structure and function.

#### Architecture of RNADiffFold

As depicted in Figure 2A, RNADiffFold consists of the diffusion model component and the condition construction unit. In the left branch, the input RNA sequence undergoes a conditional construction unit to obtain four types of feature representations: one-hot encoding, probability maps, embeddings from RNA-FM, and attention maps from RNA-FM. In the right branch, the RNA secondary structure is represented as an *L × L* binary contact map. During the diffusion process, discrete noise is gradually injected to disrupt the original contact map, and after *T* time steps, the contact map transitions into completely random noise. In the reverse diffusion process, denoising is performed using a U-Net denoising network, combined with sequence features outputted by the conditional control unit, to progressively restore the original contact map. Once the model training is completed, given a randomly sampled noise *x_T_* and an RNA sequence, the progressive denoising process can predict the secondary structure contact map.

**Fig. 2.**
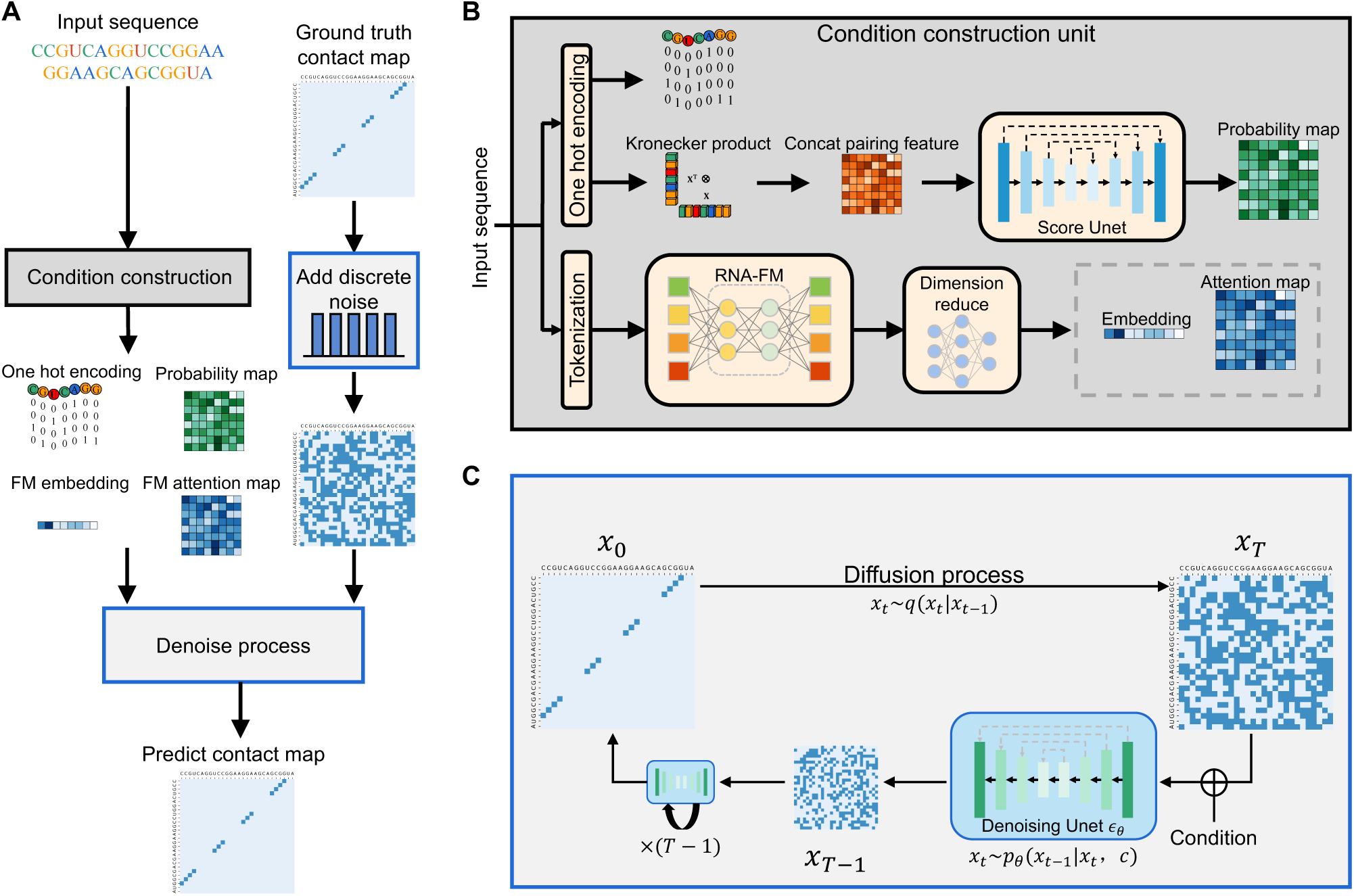
The Pipeline of RNADiffFold. (**A**) Overview of RNADiffFold workflow as a supervised task. Given the input sequence, we construct the representation through the condition construction stage. During the training stage, discrete noise is incrementally added to the ground truth contact map. Then, we gradually denoise the map conditioned by the sequence representation. In the prediction stage, given the sequence and a map randomly sampled from the categorical distribution, we generate candidate maps with different seeds and vote for the most reasonable one. (**B**) Details of condition construction. We leverage RNA-FM and UFold score networks with their respective pre-trained weights. Following the corresponding operations, we get four condition representations: one hot encoding, probability map, FM embedding, and FM attention map. (**C**) Details of the diffusion process. We leverage multinomial diffusion and a Unet network as the denoising network for training and inferring.

#### Diffusion process

As illustrated in Figure 2C, RNADiffFold employs a shared-weight neural network to learn the progressive reconstruction of data over *T* steps. The diffusion process of RNADiffFold is implemented based on multinomial diffusion [26], but there are some differences. Specifically, the model deals with binary contact maps where pixel values are limited to two representations: 0 and 1. The initial *x*_0_ is a *L × L* tensor with deterministic 0-1 relationships, where *L* represents the sequence length. To undergo denoising learning via U-Net, each pixel is embedded into an 8-dimensional vector, resulting in a *L × L ×* 8 tensor representation. Subsequently, using the Gumbel-Softmax method [35], discrete noise is gradually added to the sample at each timestep *t* through the forward process defined as follows:

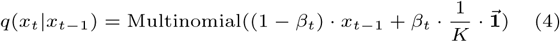

where *x_t_* represents a contact matrix at time *t* in one-hot encoded format *x^ij^ ∈ {*0, 1*}^K^* (*K* = 2, 0 *≤ i, j ≤ L*). *⃗***1** is an all-one matrix. *β_t_* is the chance of resampling another pairing possibility uniformly. We apply the cosine schedule [31] to avoid spending many steps on high-noise problems. As *t* approaches *T*, *β_t_* is adjusted to approximate 1, making the distribution closer to the uniform distribution. Since it is Markovian, we can sample arbitrary *x_t_* directly based on *x*_0_ as:

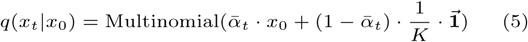

where *α_t_* = 1 *− β_t_*, and *α*^-^*_t_* = Π*^t^_τ=1_* α_τ_. Using Eq. 4 and Eq. 5 we can derive the categorical posterior *q*(*x_t−_*_1_*|x_t_, x*_0_) in the following form:

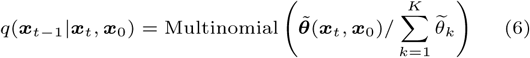

where 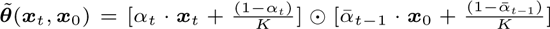. The generative diffusion process is defined as:

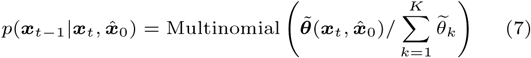

where *x*^_0_ = *µ*(*x_t_, t, c*) is a neural network to predict the initial contact matrix *x*^_0_ given the previous step state *x_t_* and condition *c ∈ {c_onehot_, c_u_, c_emb_, c_attn_}*. Note that, we parametrize *p*(*x_t−_*_1_*|x_t_*) using the probability vector from *q*(*x_t−_*_1_*|x_t_, x*^_0_). The main difference between these two processes lies in the fact that the forward diffusion process introduces to the data, making it independent of data or condition. Conversely, the generative diffusion process relies on the provided condition and a comprehensive observation of the preceding step.

The training objective of this diffusion process is to minimize the expected Kullback–Leibler(KL) divergence between Eq. 7 and Eq. 6, following a similar form as in Eq. 3:

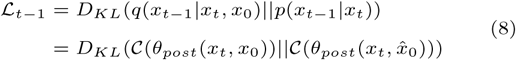

Furthermore, *L_T_* corresponds to the second term in Eq 3. Given that *x*_0_ is one-hot encoded, the computation of *L*_0_ can be expressed as:

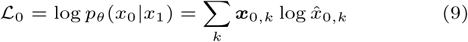

Details of denoising the Unet network are illustrated in Supplementary Figure S1. Benefiting from the flexibility of Unet to handle inputs of different sizes, we can also process variable-length sequence data.

#### Condition construction unit

The condition construction is depicted in Figure 2B. To incorporate features of the input sequence into the reverse diffusion process, the most intuitive strategy is to perform one-hot encoding on the sequence. Experimental results demonstrate that when using one-hot encoded features *c_onehot_* as conditions, RNADiffFold exhibits a certain level of prediction capability, comparable to state-of-the-art methods on some datasets. However, its predictive performance decreases when facing more complex scenarios. Therefore, it is necessary to construct additional neural networks to process input sequences, thereby extracting more meaningful features and generating sequence representations containing rich additional information.

As illustrated in Figure 2B, we adopt two neural networks as feature extractors, namely the Ufold scoring network and the pre-trained RNA-FM model. Here, the conditional representations from Ufold consist of a probability matrix *c_u_*, while those from RNA-FM consist of a sequence embedding *c_emb_*, and an attention map *c_attn_*. Given an RNA sequence of length L, denoted as **s** = (*s*_1_*, s*_2_*, …, s_L_*), where each *s_i_ ∈ {A, U, C, G, N}* and *N* represents an unknown state, the computation methods for the four types of feature representations are described as follows:

*c_onehot_* is the sequence feature from one-hot encoding, *c_onehot_ ∈ {*0, 1*}*^4*×L*^. The encoding rules are as follows: *A*: (1, 0, 0, 0), *U*: (0, 1, 0, 0), *C*: (0, 0, 1, 0), *G*: (0, 0, 0, 1), *N*: (0, 0, 0, 0).

*c_u_* is the probability matrix outputted by the scoring network from Ufold. Specifically, firstly, the Kronecker product is computed between *c_onehot_* and itself, and then the dimensions are adjusted to transform *c_onehot_* into a tensor *c_kronecker_ ∈ {*0, 1*}*^16*×L×L*^. Subsequently, to address the sparsity issue in class-like representations, *c_kronecker_* is concatenated with an additional pairing probability matrix used in CDPFold [36], resulting in a tensor *c_input_* of size 17 *×L×L*. Thus, the obtained feature representation considers all potential pairing possibilities without imposing explicit constraints, thereby enabling the prediction of more complex structures. Finally, *c_input_* is inputted into a U-Net network to produce the probability tensor *c_u_*, with dimensions of 8 *× L × L*. In this process, to meet the dimension requirements of the diffusion process, the last layer of the U-Net network is adjusted so that its output dimension is exactly 8. It is noteworthy that apart from the last layer, the initial weights of the U-Net are sourced from the pre-trained Ufold model. Subsequently, fine-tuning operations are conducted to ensure the entire network performs optimally for the current task.

*c_emb_* and *c_attn_* are respectively the one-dimensional sequence embedding and the two-dimensional attention map from RNA-FM. Specifically, the RNA sequence is inputted into the pre-trained RNA-FM model. From each encoder block’s Multi-Head Attention (MHA) layer, attention maps of dimensions 20 *×L×L* are obtained, and from the output layer, the sequence embedding *c_emb_* of dimensions 640 *×L* are derived, where 640 represents the embedding dimension, and 20 represents the number of attention heads. Subsequently, the attention maps from the 12 encoder blocks are integrated to form the final attention map *c_attn_* of dimensions 240 *× L × L*. *c_emb_* contains meaningful biological information. As argued in the [34], the multiple attention heads in different layers of the Transformer encoder module should theoretically capture structural information by focusing on the strength of correlations between different positions in the input sequence. Therefore, *c_attn_* contains information related to the structure.

#### Datasets preparation

We evaluated RNADiffFold using several datasets for both within-family and cross-family scenarios, following prior work [19, 21, 22]. These datasets include RNAStrAlign [37], ArchiveII [38], bpRNA-1m [39], bpRNA-new [20], and PDB. Specifically, the RNAStrAlign, ArchiveII, and bpRNA-1m datasets were utilized for evaluating within-family performance, while the bpRNA-new and PDB datasets were employed for assessing cross-family performance.

RNAStrAlign comprises 30,451 unique sequences distributed among 8 RNA families, whereas ArchiveII consists of 3,975 sequences spanning 10 RNA families. Due to their similar family distributions, RNAStrAlign is often used as a training set and ArchiveII as a test set. The bpRNA-1m dataset contains 102,318 sequences from 2,588 families and is considered one of the most exhaustive datasets used for benchmarking RNA secondary structure prediction methods. On the other hand, the bpRNA-new dataset was derived from Rfam14.2 [40] and includes 1,500 new RNA families detected by newly developed techniques [41]. Furthermore, the PDB dataset, collected from the Protein Data Bank [42], comprises high-resolution RNA structures with resolutions less than 3.5°*A* (as of April 9, 2020).

Before using the datasets, we performed preprocessing steps to enhance computational efficiency and reduce redundancy. Specifically, for the RNAStrAlign dataset, we removed sequences longer than 640 nucleotides and randomly split the dataset into training, validation, and test sets in an 8:1:1 ratio. Redundant sequences were removed using methods from E2Efold [18] and Ufold [19]. For the ArchiveII dataset, which was used solely for testing, a similar strategy as the RNAStrAlign dataset was adopted to reduce redundancy. Regarding the bpRNA-1m dataset, We used CD-HIT-EST [43] to remove sequences with over 80% similarity, following the methods of MXfold2 [20] and SPOT-RNA [21]. The remaining data, referred to as bpRNA, was divided into training, validation, and test sets labeled TR0, VL0, and TS0, respectively. For the bpRNA-new dataset, sequences with over 80% similarity were removed using CD-HIT-EST, and sequences longer than 500 nucleotides were filtered out. For the PDB dataset, we applied the same dataset division as SPOT-RNA2 [22], resulting in training (TR1), validation (VL1), and various test sets (TS1, TS2, TS3, TS-hard).

Furthermore, we analyzed the relationship between sequence length and the number of base pairs in each dataset, finding a linear relationship for most samples, as shown in Supplementary Figure S2. However, some samples deviated from the fitted line, with most of the deviations related to sequences containing low pairing information and some being completely unpaired. Thus, sequences longer than 200 nucleotides with fewer than 10 base pairs were removed from the training set to reduce the impact of low-pairing sequences. Validation and test sets remained consistent with previous work. The final dataset used in this study was slightly smaller than that used in previous studies.

To enhance the transfer ability of the model in handling unknown RNA families, we employed data augmentation similar to Ufold [19]. In the bpRNA-new dataset, 20% *−* 30% of nucleotides were randomly mutated to create new data. Then, sequences with over 80% similarity to real sequences were removed using CD-HIT-EST, yielding 2717 mutated sequences (named Mutate-seq) for supplementary training. Labels for Mutate-seq were generated using Contrafold [44], a probabilistic method extending stochastic context-free grammars (SCFGs) with discriminative training objectives and flexible feature representations.

In summary, the dataset configuration for this study includes multiple training sets derived from RNAStrAlign, bpRNA TR0, and Mutate-seq, along with TR1 from the PDB dataset utilized for cross-family fine-tuning. For within-family evaluations, the test sets include sequences from ArchiveII and bpRNA TS0. Conversely, cross-family testing employs bpRNA-new and multiple test sets from PDB (TS1, TS2, TS3, and TS-hard). The detailed statistics of the final datasets employed for the training, validation, and testing phases are systematically presented in Supplementary Table S1.

We also observed that approximately 80% of the sample sequences in the original data were concentrated below 160 nucleotides in length (23,374 sequences between 30 and 160 nucleotides, 3,202 sequences between 160 and 320 nucleotides, and 3,119 sequences between 320 and 640 nucleotides), as shown in Figure S3A, which hindered the model’s ability to learn long-range RNA interactions. To address this issue, a data balancing strategy [45] was employed. Specifically, sequences with lengths between 160 and 640 nucleotides were selectively duplicated in the training set to increase their representation in the dataset. After duplication, the number of sequences with lengths between 160 and 640 nucleotides increased to 48658. This initiative effectively mitigated the impact of data imbalance on model training. Figure S3B shows the distribution of sequence length and number of base pairs after data balance. In addition, to improve the training efficiency, we also developed a data bucketing strategy to process the above datasets, and the details are given in Supplementary Section 1.1.

## Experiments and results

### Evaluation criteria and experimental setup

To comprehensively evaluate the prediction accuracy of RNADiffFold, we employed precision (*Prec*), recall (*Recall*), and F1 score as evaluation metrics, defined as follows:

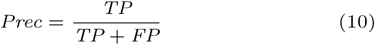

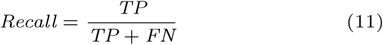

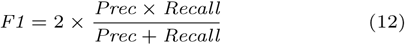

where *TP* represents the number of correctly predicted base pairs (true positives), *FP* represents the number of incorrectly predicted base pairs (false positives), and *FN* represents the number of base pairs in the reference structure that were not predicted (false negatives).

RNADiffFold is implemented using Pytorch [46]. We set the number of diffusion steps *T* as 20 during the diffusion process, considering that the contact map contains sparse classification information. A large *T*, on the contrary, tends to degrade prediction performance. Adam [47] is employed as the optimizer with a learning rate of 1e-3. The model undergoes training for a maximum of 400 epochs, with validation performed every 20 epochs. Early stopping with a patience of 5 is implemented to prevent overfitting. Supplementary Table S10 details all the hyper-parameters of the experiment.

The training process is divided into two stages. Firstly, the model is trained on the RNAStrAlign training set, TR0, and Mutate-data, followed by testing on ArchiveII, TS0, and bpRNA-new to evaluate its generalization ability. Secondly, using the model weights obtained in the first stage, fine-tuning is performed on the PDB dataset, and testing is conducted on TS1, TS2, TS3, and the challenging TS-hard dataset to validate the model’s performance in different scenarios. All experiments are carried out on a single Nvidia A40 GPU and an Intel(R) Xeon(R) Gold 6326 CPU @ 2.90GHz. An analysis of computational efficiency is provided in Supplementary Section 1.3 and Table S12.

### Baselines

We compared RNADiffFold with ten competitive baseline methods for RNA secondary structure prediction, including six energy-based approaches and four deep learning-based methods. Among the energy-based methods, Mfold [48] utilizes nearest-neighbor energy parameters and dynamic programming to find the structure with the lowest energy. Linearfold [13] combines dynamic programming with beam search to enhance efficiency. RNAfold [49] integrates dynamic programming algorithms with a thermodynamic-based energy model to predict optimal RNA structures. RNAStructure [11] calculates minimum free energy and optimizes prediction results based on experimental data. Contrafold [44] uses conditional log-linear models, extending to SCFGs, and integrates thermodynamic parameters. ContextFold [50] introduces a fine-grained model with a large parameter set (approximately 70,000).

For the deep learning-based methods, SPOT-RNA [21] employs a CNN and BiLSTM architecture and utilizes transfer learning to enhance performance. MXFold2 [20] combines deep learning with thermodynamic parameters for more accurate predictions. E2Efold [18] innovatively integrates the Transformer model with an unrolling algorithm, imposing hard constraints on RNA secondary structure. UFold [19] utilizes U-Net and a structure similar to image inputs to capture long-range dependencies effectively. Due to the lack of reproducible results from SPOT-RNA and ContextFold in experiments, we referenced the experimental data of UFold for comparison analysis.

### Performance on within-family datasets

The “within-family” datasets indicate that the RNA families in the test set are highly similar to those in the training set. As reported in previous studies, evaluation results on these datasets effectively reflect the model’s predictive performance. As shown in Figure 3 and Supplementary Table S2, for the ArchiveII test set, RNADiffFold demonstrates good predictive performance with an F1 score of 0.880, surpassing all energy-based methods and ranking among the top in deep learning-based methods. Its F1 score is slightly lower than UFold and RNA-FM, possibly due to information loss from the conditional construction strategy. Since ArchiveII contains a limited number of families and species, RNADiffFold achieves comparable predictive performance using only one-hot encoding as the condition, as further confirmed in subsequent ablation experiments.

**Fig. 3.**
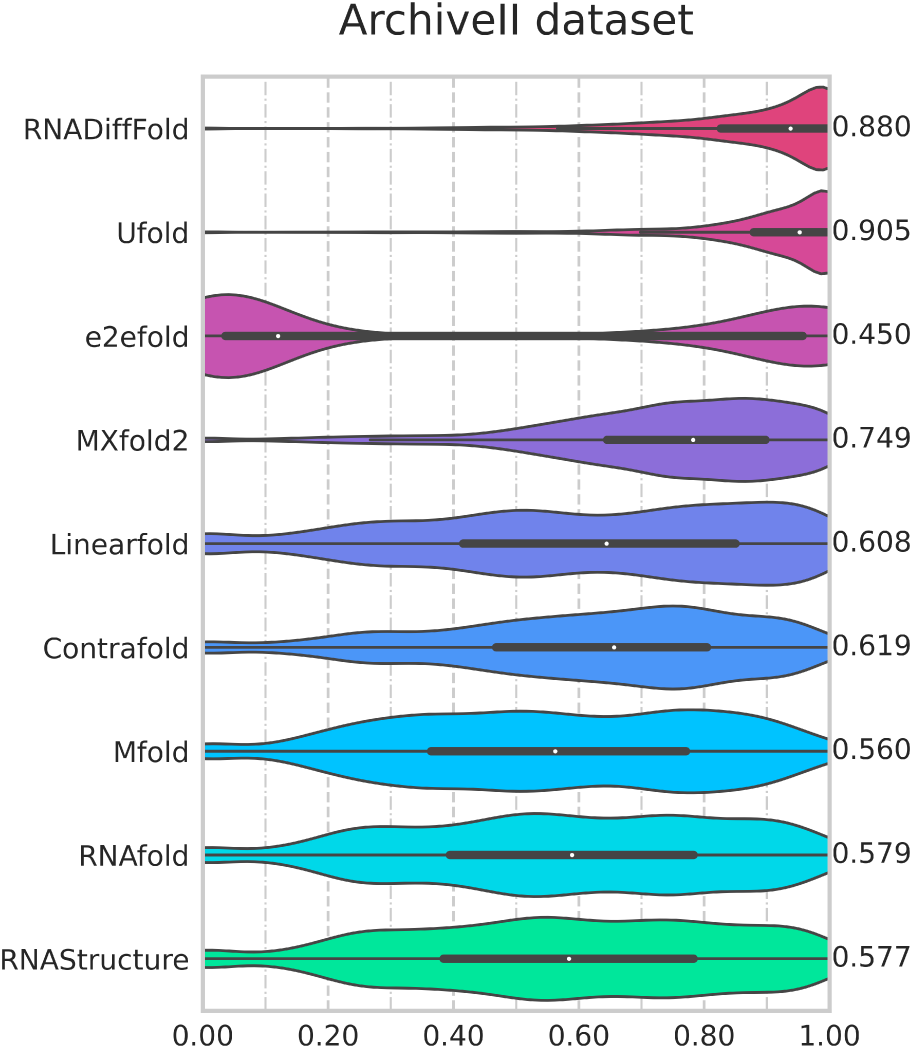
Violin plot on the ArchiveII dataset. Visualization of F1 value of RNADiffFold against other methods.

In contrast, the bpRNA dataset from Rfam12.2 [51] encompasses a broader range of RNA families. Evaluation results on its test set, TS0, show RNADiffFold achieves an average F1 score of 0.711, surpassing all energy-based and deep learning-based methods (Figure 4 and Supplementary Table S2). Specifically, RNADiffFold shows a notable 8.7% improvement in F1 score compared to UFold and a 2.7% increase compared to RNA-FM. This significant performance enhancement highlights the superiority of RNADiffFold in the field of RNA secondary structure prediction, demonstrating that its outstanding performance results from its unique algorithm design and feature fusion strategy, rather than a simple combination of the UFold scoring network and RNA-FM output features.

**Fig. 4.**
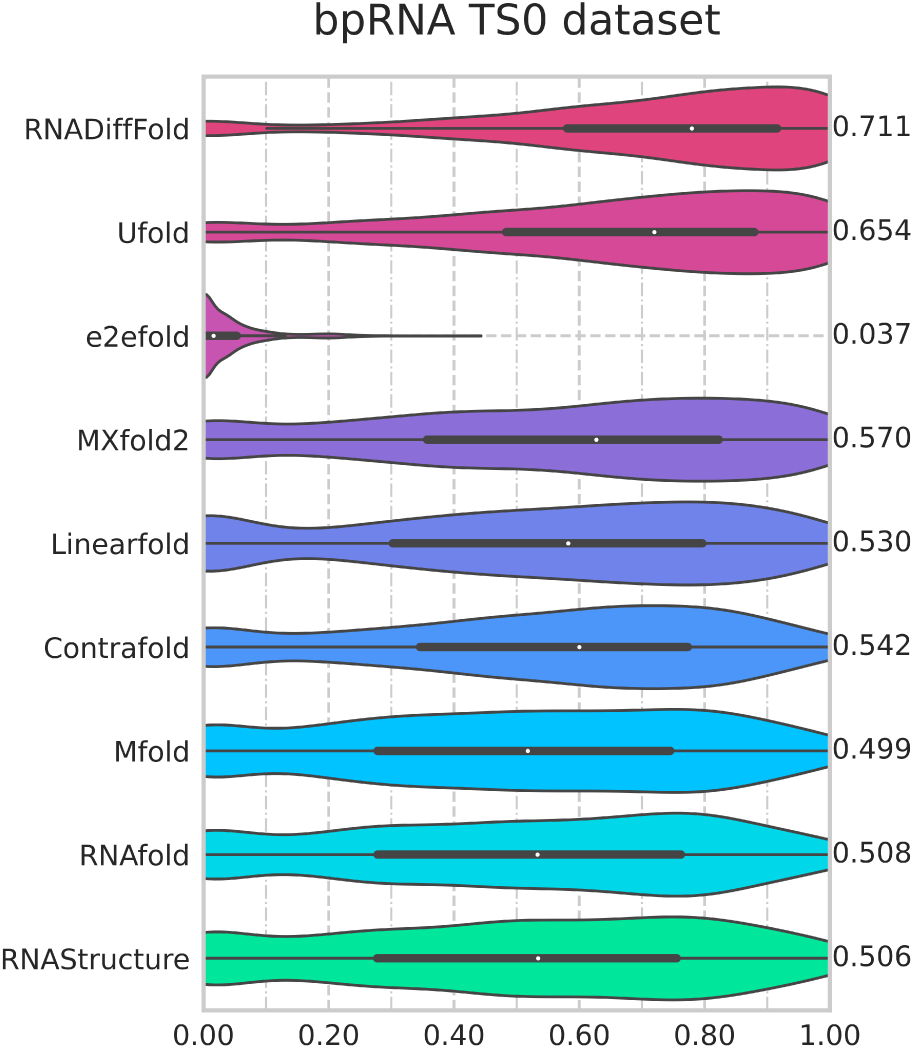
Violin plot on the bpRNA TS0 dataset. Visualization of F1 value of RNADiffFold against other methods.

To comprehensively assess RNADiffFold’s capability in predicting long-range base pairs, we followed the methodology of UFold [19] and conducted an in-depth analysis on the bpRNA TS0 dataset. For each RNA sequence of length *L*, we classified base pairs and non-base pairs with intervals exceeding *L/*2 as long-range base pairs. After rigorous screening, we selected 993 samples from a total of 1,304 that met the criteria for precision, recall, and F1 score calculations. Comparing RNADiffFold with other mainstream methods, the results, presented in Table 1 and Figure S4, demonstrate RNADiffFold’s outstanding performance on long-range base pair RNA data. Compared to UFold, RNADiffFold’s predicted precision and recall are closer, indicating a more stable predictive performance. This stability likely stems from the effective integration of the pre-trained UFold scoring network with RNA-FM output features, providing RNADiffFold with rich contextual and structural information, thereby enhancing its accuracy in predicting long-range base pairs.

**Table 1.** F1 scores of RNADiffFold compared with other learning-based methods on the TS0 dataset of long-range base pairing.

| Method | Precision | Recall | F1 |
| --- | --- | --- | --- |
| RNADiffFold | **0.748** | 0.764 | **0.739** |
| Ufold | 0.644 | **0.765** | 0.687 |
| SPOT-RNA<sup>a</sup> | 0.361 | 0.492 | 0.403 |
| MXfold2 | 0.318 | 0.450 | 0.360 |
| e2efold | 0.038 | 0.084 | 0.043 |

<sup>a</sup> The results of SPOT-RNA are cited from Ufold[19]

### Performance on cross-family datasets

The “cross-family” datasets indicate that the RNA family species in the test set are entirely distinct from those in the training set. Performance on such datasets, particularly for deep learning-based methods, better reflects their understanding of RNA folding patterns and their ability to generalize to unknown families. To enhance the model’s generalization performance, we adopted UFold’s data augmentation strategy, generating 2,717 mutated sequences with mutation rates between 20% and 30% from the original sequences. These sequences were predicted for their secondary structures by Contrafold, serving as pseudo-labels. These mutated datasets were then integrated into RNAStrAlign and TR0 for training and comprehensively evaluated on three different test sets.

As shown in Figure 5 and Supplementary Table S3, RNADiffFold outperformed other methods, demonstrating its superior performance in predicting the secondary structures of RNA sequences from unknown families. Unlike baseline methods like MXfold2, which incorporates thermodynamic constraints, or UFold, which employs biological-based hard constraints, RNADiffFold predicts the secondary structures of new, unknown family sequences by learning the distribution of RNA conformations. Additionally, we analyzed the performance across different sequence lengths by partitioning the bpRNA-new dataset at 100 nt intervals. As shown in Figure S5, the performance of most methods deteriorates with increasing length. RNADiffFold maintains excellent predictive performance for sequences below 300 nt but decreases more for sequences between 300 and 500 nt, possibly due to the absence of constraints.

**Fig. 5.**
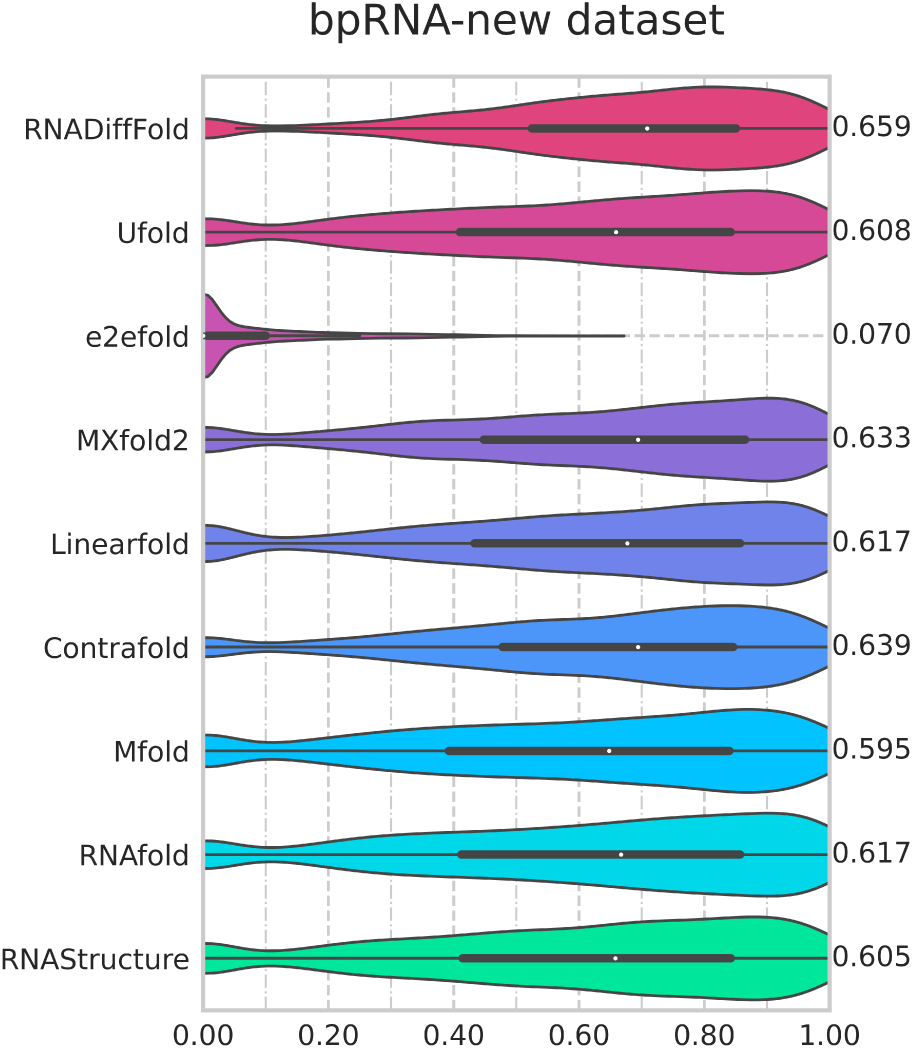
Violin plot on the bpRNA-new dataset. Visualization of F1 value of RNADiffFold against other methods.

To comprehensively evaluate RNADiffFold in cross-family scenarios, we conducted additional validation using the PDB dataset. The PDB dataset [42] contains secondary structure information extracted from high-resolution RNA 3D structures, providing a reliable basis for evaluation. Following the same dataset partitioning method as SPOT-RNA2 [22], we fine-tuned the pre-trained model on the PDB dataset. Figure 6 shows that RNADiffFold achieved an average F1 score of 0.736, demonstrating comparable performance with other methods. Further analysis on different test sets within PDB (TS1, TS2, and TS3) is detailed in Table S4 and Figure S6. We also constructed a more challenging test set, TS-hard, which was curated from TS1 and TS3 by excluding similar sequences using BLAST-N [52] and INFERNAL [53] models. The results on TS-hard, shown in Table S5 and Figure S7, validate that RNADiffFold maintains comparable performance with most methods even in challenging prediction tasks.

**Fig. 6.**
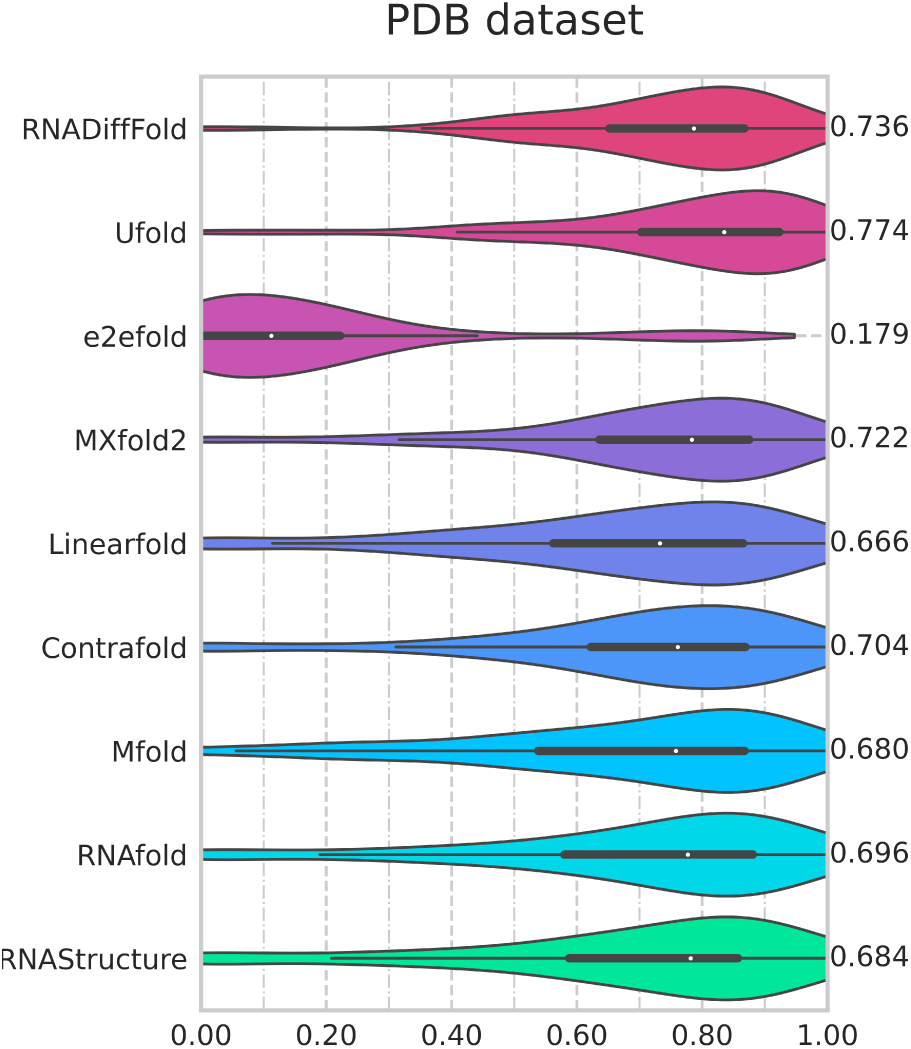
Violin plot on the PDB test dataset encompassing TS1, TS2 and TS3. Visualization of F1 value of RNADiffFold against other methods.

To further substantiate RNADiffFold’s predictive capability, we employed two statistical significance assessment methods: an individual t-test-based method and a bootstrapping-based method. The results in Table S6 indicate that RNADiffFold significantly outperforms other methods statistically, with most p-values being less than 0.05. For the relatively small-scale PDB dataset, bootstrapping proved to be a more effective evaluation method. Table S7 and Figure S8 present the 95% confidence intervals of F1 scores for all compared methods on the PDB test sets (TS1, TS2, TS3), providing robust support for our conclusions.

### Multiple sampling and voting strategy testing experiments

To further enhance the model’s performance, we implemented a strategy based on multiple sampling and voting. Given that RNADiffFold is a generative prediction method capable of learning the distribution of secondary structures, we used random seeds to generate a set of predicted structures. We then conducted cluster analysis by calculating the similarity between these predictions and selected the most frequently occurring structure from the largest cluster as the final prediction. When no clear clusters were present, a random selection method determined the final prediction. This strategy resulted in a performance improvement, as shown in Table S8. By increasing the number of samples from 1 to 10, moderate improvements were observed in the evaluation metrics across the four test sets. This outcome suggests that the multiple sampling and voting strategy helps reduce the uncertainty of model predictions, thereby enhancing the accuracy of RNA secondary structure prediction. The performance shown in Figures 3 to 6 all adopted this strategy.

### Visualization

To intuitively demonstrate the model’s ability to capture the details of RNA secondary structures, we randomly selected RNA sequences from different RNA family species and visually compared the predicted structures from RNADiffFold with those from two other top-performing methods, Ufold and MXfold2, as well as the ground truth. The predicted results were converted into .ct format based on base pairing relationships and visualized using VARNA [54]. Figure 7 shows that the structures predicted by RNADiffFold are closer to the ground truth than those predicted by other methods, demonstrating improved accuracy in detail.

**Fig. 7.**
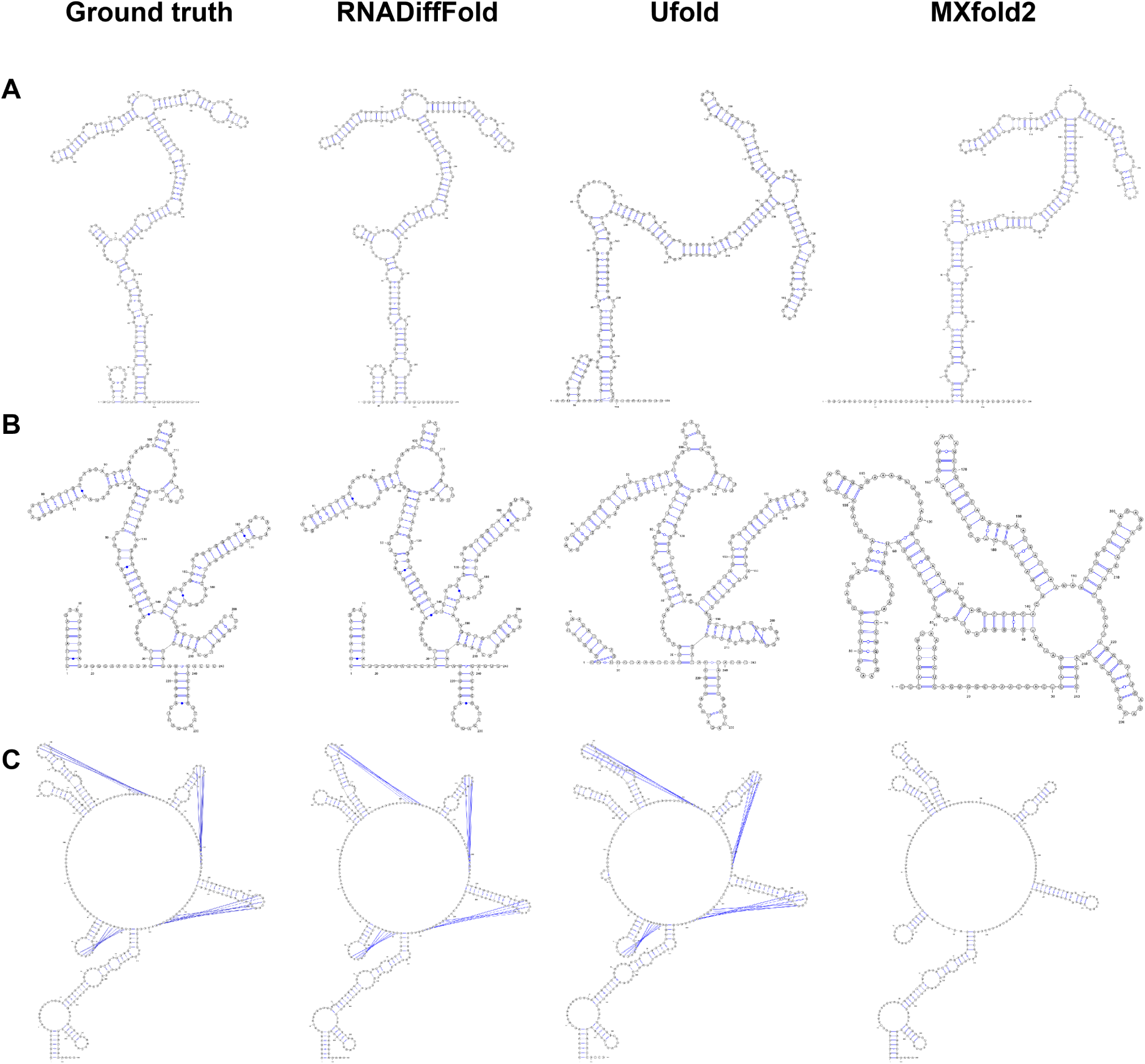
Visualization compares three examples predicted by RNADiffFold with those from two other methods against the ground truth. The RNA sequences are from the following families: (**A**) Aspergillus fumigatus, recorded in the SRPDB database; (**B**) Alphaproteobacteria subfamily 16S rRNA, recorded in the RNAStrAlign database; and (**C**) Escherichia coli, recorded in the tmRNA database. The results indicate that RNADiffFold aligns more closely with the ground truth in detail than the other two methods.

We also validated RNADiffFold’s ability to capture dynamic structural features using CoDNaS-RNA [55], a database containing diverse RNA conformational ensembles. Considering data quality and sequence length, we selected 14 clusters, each containing two different conformations of the same sequence. Among these, 9 clusters had extremely similar sequences, differing by only 1-2 bases. The experimental results indicate that RNADiffFold successfully predicted the structural profiles of 11 clusters and captured the different conformations to a certain extent. Figure S10 shows examples of 4 successfully predicted clusters, of which Cluster 189 represents the prediction of 9 similar clusters. The failed cases occurred because the ground truth sequences tended to be completely unpaired or contained pairings that violated standard pairing rules, making accurate predictions challenging, as shown in Figure S11.

### Ablation Study

In this section, we explore the impact of different conditioning strategies on the performance of RNADiffFold while maintaining other experimental parameters constant. The versions with different conditions are labeled as follows: v1: using only the one-hot encoding *c_onehot_* of the given sequence; v2: utilizing the probability map *c_u_* from the Ufold scoring network; v3: employing both *c_onehot_*, *c_emb_*and *c_attn_* from RNA-FM simultaneously; v4: utilizing *c_onehot_*, *c_u_*, and *c_emb_* as conditions simultaneously; v5: RNADiffFold’s final form, incorporating all four conditions simultaneously.

Figure 8 and Table S9 present the comprehensive performance evaluation results of RNADiffFold on four test sets. Key observations drawn from this study are as follows: (1) On the ArchiveII test set and TS0, RNADiffFold exhibits competitiveness with state-of-the-art methods (Ufold and RNA-FM) when using only *c_onehot_* as a condition (version v1). However, the performance of the v1 is relatively limited when faced with sequences from unknown families, indicating the inadequacy of one-hot encoding in feature extraction. (2) The probability map *c_u_* from the Ufold scoring network (version v2) contributes more significantly to cross-family data, while *c_emb_* and *c_attn_* from RNA-FM (version v3) play a more significant role in within-family data. Integrating these output features as sequence condition information achieves more balanced performance on both within-family and cross-family data. (3) Comparing versions v4 and v5 highlights the utility of the attention map *c_attn_*. Introducing *c_attn_* results in slight performance improvements on almost all datasets, albeit modest. This may be due to partial feature information loss from excessive dimensionality reduction. To determine if the exceptional performance of RNADiffFold primarily stems from its diffusion process, we conducted further comparative experiments detailed in Supplementary Section 1.2 and Table S11. The results confirm that the integration of features from Ufold and RNA-FM alone does not account for the superior outcomes observed; the diffusion process is pivotal to its success. Additionally, early experiments explored the relationship between different diffusion step sizes and model performance. Unexpectedly, smaller step sizes performed better in the diffusion process, as shown in Figure S9. This may be because the secondary structure contact map contains sparser information compared to real-world images. Conversely, larger step sizes may make it difficult for the model to explore the conformation distribution space. These findings provide valuable guidance for further optimizing RNADiffFold.

**Fig. 8.**
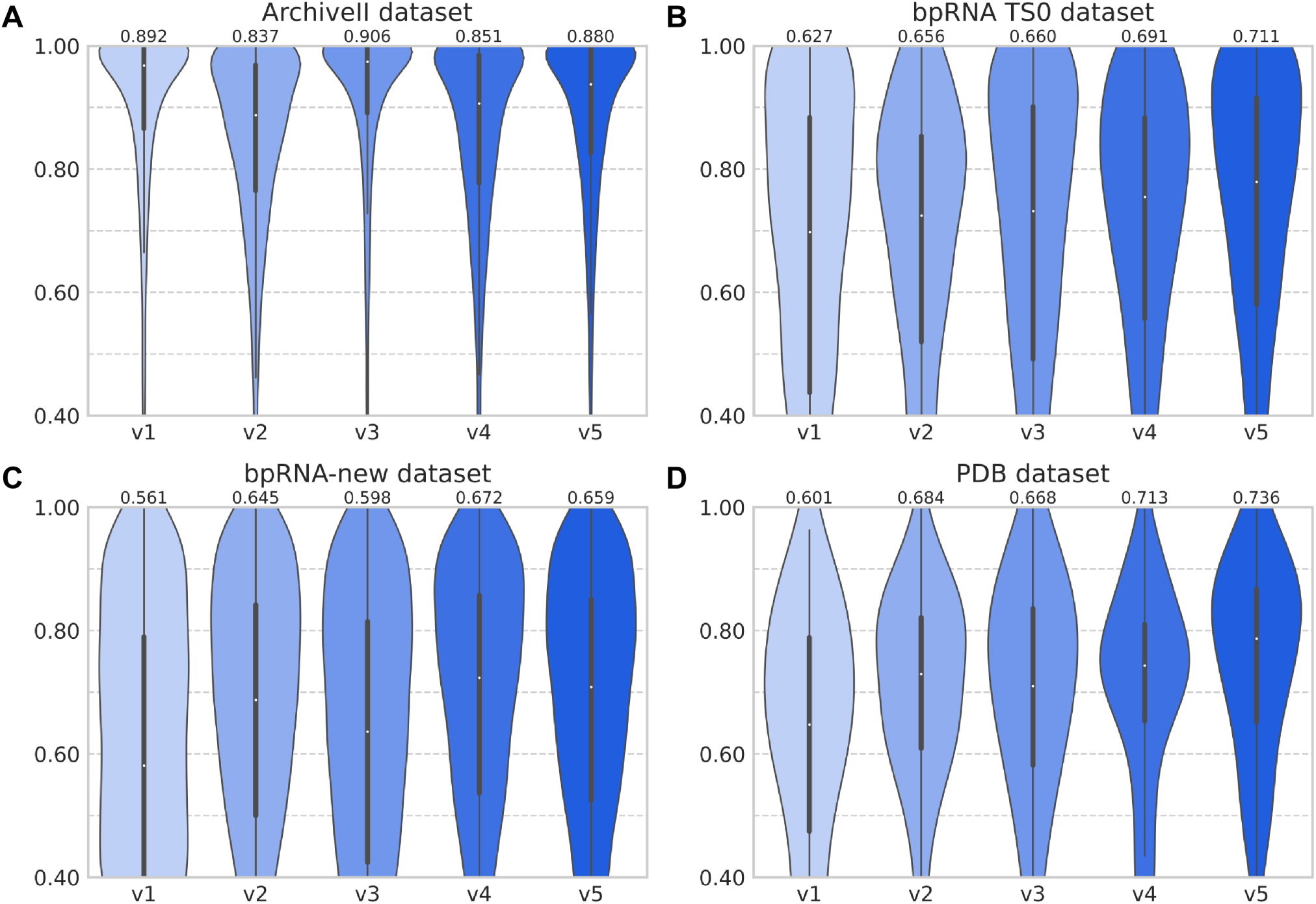
F1 values on the four datasets. (within-family datasets ArchiveII (**A**), bpRNA TS0 (**B**) and cross-family datasets bpRNA-new (**C**), PDB datasets (**D**)). v1-v5 are different condition construction strategies combined with the diffusion process. v1: one-hot encoding *c_onehot_*; v2: probability map from the Ufold score network *c_u_*; v3: one-hot encoding *c_onehot_*, output from RNA-FM *c_emb_, c_attn_*; v4: one-hot encoding *c_onehot_*, probability map from the Ufold score network *c_u_*, output from RNA-FM *c_emb_*; v5: full version of RNADiffFold.

## Discussion

In this paper, we present RNADiffFold, a novel RNA secondary structure prediction method based on the discrete diffusion model. Unlike traditional deep learning-based prediction methods, RNADiffFold treats secondary structure prediction as a pixel-level contact map segmentation task, utilizing discrete diffusion processes to capture conformational distributions. During the forward diffusion process, noise following a uniform distribution is gradually injected into the true contact map until it is completely randomized. In the reverse process, RNADiffFold employs a UNet as the denoising network. Additionally, we propose an effective conditioning strategy to extract condition information from RNA sequences to guide structure prediction.

RNADiffFold offers several significant advantages over previous methods. First, traditional approaches typically predict secondary structures in a deterministic manner, which contradicts the dynamic folding nature of RNA. RNADiffFold, however, explores the conformational space of RNA through diffusion processes without imposing any explicit hard constraints, allowing it to predict non-canonical pairings arising from tertiary interactions. Second, our proposed conditioning strategy leverages the strengths of different models. From another perspective, RNADiffFold can be viewed as a downstream task of pre-trained models, avoiding the need for embedding complex prior knowledge. Experimental results demonstrate that RNADiffFold achieves competitive performance in predicting RNA secondary structures compared to current deep learning and energy-based methods. Additionally, the method exhibits the ability to capture dynamic RNA features. Further experiments on intra-family and inter-family datasets validate the effectiveness and robustness of RNADiffFold.

Although RNADiffFold demonstrates outstanding predictive performance, there is still room for improvement. First, a more specific design of the loss function for training the diffusion model is needed to address contact map features containing sparse information. Unlike real-world images, contact maps contain less category information, with most regions classified as “0” class. Therefore, resources should be reduced in these easily learnable regions. Second, adopting more efficient conditional pre-training models could enhance performance. Due to computational cost limitations, this study has not retrained a condition control model tailored for the diffusion model. Theoretically, using more suitable and thoroughly trained pre-training models to extract sequence features as conditions is expected to further improve the performance of RNADiffFold.

## Supporting information

Supplementary Information for RNADiffFold

## Key Points

- We introduce RNADiffFold, a novel discrete diffusion framework that treats RNA secondary structure prediction as a pixel-level segmentation task. This approach effectively captures the distribution of secondary conformations, unveiling valuable biological insights embedded in the primary sequence.
- A condition construction strategy is proposed to construct sequence features, enabling flexible feature design to enhance prediction performance or cater to specific tasks.
- Experiments show that RNADiffFold surpasses previous energy-based and recent learning-based methods on within- and cross-family datasets, demonstrating the effectiveness and robustness of RNADiffFold.

## Data and Code Availability

The code to reproduce our experiments and source data is available at https://github.com/HIM-AIM/RNADiffFold under an MIT License.

## Author Contribution

Z.W., Y.F., P.Y., and X.L. conceived the research project. X.L. and Y.P. supervised and advised the research project. Z.W. and Y.F. designed and implemented the RNADiffFold framework. Z.W., Y.F., Q.T., and Z.L. conducted the computational analyses. Z.W., Y.F., and X.L. wrote the manuscript. All the authors discussed the experimental results and commented on the manuscript.

## Acknowledgments

We acknowledge helpful discussions with members of the AIM lab. The authors thank the anonymous reviewers for their valuable suggestions.

## Funding

This work is supported in part by funds from the National Key Research and Development Program of China (2022YFC3600902).

## Competing Interests Statement

The authors declare no competing interests.

## Notes

### Competing Interest Statement

The authors have declared no competing interest.

### Summary of Updates

The author signature order and affiliations have been updated; article details have been updated; supplementary files have been updated.

