## Supplementary Information for RNADiffFold for "RNADiffFold: Generative RNA Secondary Structure Prediction using Discrete Diffusion Models"

Oct 2024

### 1 Supplementary Sections

#### 1.1 Data bucketing strategy

Previous approaches typically padded all training data to the maximum sequence length for batch training, which resulted in lower training efficiency. To improve training efficiency, we implemented a data bucketing strategy to handle the datasets. Specifically, for each dataset, sequences were sorted by length and then binned with intervals of 80 nucleotides, ensuring that sequences within the same bucket had similar lengths. Subsequently, sequences within each bucket were padded to the maximum length of that bucket. For example, sequences with lengths between 80 to 160 nucleotides were grouped into the same bucket and uniformly padded to 160 nucleotides. This approach ensured that shorter sequences were not padded to excessively large lengths. During training, buckets were utilized for training, and the batch size was set to 1. This bucketing strategy alleviated unnecessary computations in padding regions, ensuring efficient data processing and reducing training time by approximately 50%.

#### 1.2 Additional ablation experiments

To further explore whether the exceptional performance of RNADiffFold is mainly attributable to the diffusion process, we conducted additional comparative experiments:

- Simple Combination (Combination_1): We directly combined the output probability maps from Ufold and RNA-FM, processed through the post-processing steps of Ufold without involving any trainable parameters. This setup was intended to test the effect of a straightforward combination of outputs from the two models.
- Combination with Trainable Parameters (Combination_2): To align with the parameter count of RNADiffFold, we designed a Unet network incorporating trainable parameters to merge features from the outputs of Ufold and RNA-FM.

The experimental results in Table S11 clearly show that neither the simple combination nor the combination with additional trainable parameters reached the performance level of RNADiffFold. This indicates that the superior performance of RNADiffFold cannot be achieved by merely combining the features from Ufold and RNA-FM outputs. The diffusion process plays a crucial role.

#### 1.3 Computational efficiency analysis

To evaluate the computational efficiency, we have evaluated the inference time of RNADiffFold for predicting the structure of a single RNA sequence. Detailed comparisons of inference times across various models on the RNAStralign test set are presented in Table S12. Our results indicate that the inference time for RNADiffFold is approximately 1-2 seconds per sequence on a single A40 GPU with 20 inference steps. While this performance does not show a marked advantage over other advanced methods such as Ufold, MXfold2, E2Efold, and LinearFold—mainly due to the inherently slower sampling speed of diffusion models—we observed significant efficiency when processing batch data. If the batch size is set to a large number (greater than 10), RNADiffFold's average prediction speed per sequence can compete with that of Ufold, primarily constrained by the available computer memory.

Furthermore, we have conducted a comparative analysis of the number of trainable parameters across RNADiffFold and various baseline models, which can also be found in Table S12. RNADiffFold has 836,423 trainable parameters, which only includes the diffusion model part and excludes the conditioning units, as the conditioning units utilize existing models that require no additional training. This parameter count is considerably lower compared to some models (e.g., Ufold with 8,641,377 parameters), reflecting the efficiency of RNADiffFold to some extent.

### 2 Supplementary Tables

Table S1: Datasets for our experiments consist of training, validation, and testing sets, excluding redundant sequences with low base pair ratios to the sequence length.

| Training Method | Dataset | Training set | Validation set | Test set | Test Scenario |
|---|---|---|---|---|---|
| Training | RNAStrAlign | 16184 | 2024 | / | within-family |
| | bpRNA | 10794(TR0) | 1299(VL0) | 1304(TS0) | |
| | Mutate-seq | 2717 | / | / | |
| | ArchiveII | / | / | 3911 | |
| / | bpRNA-new | / | / | 5401 | cross-family |
| Finetuning | PDB | 119(TR1) | 29(VL1) | (TS1,TS2,TS3,TS-hard) | |

Table S2: Benchmark results on the ArchiveII dataset and TS0 dataset.

| Method | ArchiveII | | | TS0 | | |
|---|---|---|---|---|---|---|
| | Prec[a] | Rec | F1 | Prec[a] | Rec | F1 |
| RNADiffFold | 0.889 | 0.878 | 0.880 | **0.717** | **0.743** | **0.711** |
| Ufold | 0.890 | 0.926 | 0.905 | 0.607 | 0.741 | 0.654 |
| e2efold | 0.467 | 0.499 | 0.450 | 0.031 | 0.073 | 0.037 |
| MXfold2 | 0.737 | 0.768 | 0.749 | 0.514 | 0.685 | 0.570 |
| Linearfold | 0.605 | 0.627 | 0.608 | 0.581 | 0.517 | 0.530 |
| Contrafold | 0.651 | 0.595 | 0.619 | 0.655 | 0.482 | 0.542 |
| Mfold | 0.579 | 0.550 | 0.560 | 0.611 | 0.452 | 0.499 |
| RNAfold | 0.613 | 0.553 | 0.579 | 0.631 | 0.446 | 0.508 |
| RNAStructure | 0.606 | 0.554 | 0.577 | 0.623 | 0.447 | 0.506 |
| RNA-FM[b] | **0.924** | **0.958** | **0.939** | 0.684 | 0.725 | 0.694 |
| SPOT-RNA[c] | 0.743 | 0.726 | 0.711 | 0.594 | 0.693 | 0.619 |
| Contextfold[c] | 0.873 | 0.821 | 0.821 | 0.529 | 0.607 | 0.546 |

[a] Pre,Rec and F1 are the macro averages of precision, recall and F1 score respectively.
[b] The results of RNA-FM is cited from RNA-FM[1].
[c] The results of SPOT-RNA and Contextfold are cited from Ufold[2].

Table S3: Benchmark results on the bpRNA-new dataset.

| Method | Prec[a] | Rec | F1 |
|---|---|---|---|
| RNADiffFold | 0.672 | 0.693 | **0.659** |
| Ufold | 0.537 | **0.723** | 0.608 |
| e2efold | 0.088 | 0.065 | 0.070 |
| MXfold2 | 0.580 | 0.718 | 0.633 |
| Linearfold | 0.645 | 0.621 | 0.617 |
| Contrafold | **0.736** | 0.578 | 0.639 |
| Mfold | 0.687 | 0.539 | 0.595 |
| RNAfold | 0.720 | 0.552 | 0.617 |
| RNAStructure | 0.704 | 0.543 | 0.605 |
| SPOT-RNA[b] | 0.635 | 0.641 | 0.620 |
| Contextfold[b] | 0.596 | 0.636 | 0.604 |

[a] Pre,Rec and F1 are the macro averages of precision, recall and F1 score respectively.
[b] The results of SPOT-RNA and Contextfold are cited from Ufold[2].

Table S4: Benchmark results on the TS1, TS2 and TS3 dataset.

| Method | TS1 | | | TS2 | | | TS3 | | |
|---|---|---|---|---|---|---|---|---|---|
| | Prec | Rec | F1 | Prec | Rec | F1 | Prec | Rec | F1 |
| RNADiffFold | 0.829 | 0.621 | 0.703 | 0.937 | 0.738 | 0.814 | 0.852 | 0.582 | 0.682 |
| Ufold | 0.781 | 0.665 | 0.712 | **0.943** | 0.848 | **0.891** | 0.849 | 0.647 | **0.731** |
| e2efold | 0.298 | 0.185 | 0.221 | 0.259 | 0.107 | 0.149 | 0.174 | 0.091 | 0.104 |
| MXfold2 | 0.788 | 0.629 | 0.695 | 0.901 | 0.708 | 0.785 | 0.811 | 0.593 | 0.679 |
| Linearfold | 0.545 | 0.731 | 0.617 | 0.696 | 0.889 | 0.770 | 0.515 | **0.812** | 0.613 |
| Contrafold | 0.603 | **0.733** | 0.657 | 0.744 | 0.878 | 0.801 | 0.578 | 0.789 | 0.655 |
| Mfold | 0.546 | 0.659 | 0.593 | 0.740 | 0.919 | 0.815 | 0.610 | 0.797 | 0.687 |
| RNAfold | 0.588 | 0.697 | 0.633 | 0.751 | **0.921** | 0.821 | 0.575 | 0.736 | 0.642 |
| RNAStructure | 0.570 | 0.687 | 0.619 | 0.742 | 0.910 | 0.813 | 0.562 | 0.731 | 0.633 |
| SPOT-RNA | **0.882** | 0.677 | **0.751** | 0.922 | 0.790 | 0.843 | **0.931** | 0.599 | 0.717 |
| Contextfold | 0.853 | 0.516 | 0.621 | 0.920 | 0.683 | 0.777 | 0.853 | 0.516 | 0.621 |

Table S5: Benchmark results on the TS hard dataset.

| Method | Prec | Rec | F1 |
|---|---|---|---|
| RNADiffFold | **0.784** | 0.541 | 0.624 |
| Ufold | 0.367 | 0.515 | 0.419 |
| e2efold | 0.099 | 0.097 | 0.093 |
| MXfold2 | 0.775 | 0.601 | **0.669** |
| Linearfold | 0.522 | 0.764 | 0.610 |
| Contrafold | 0.569 | 0.730 | 0.627 |
| Mfold | 0.573 | 0.716 | 0.630 |
| RNAfold | 0.575 | 0.732 | 0.639 |
| RNAStructure | 0.535 | 0.690 | 0.598 |
| SPOT-RNA[a] | 0.649 | **0.786** | 0.552 |
| SPOT-RNA2[a] | 0.678 | 0.731 | 0.632 |

[a] The results of SPOT-RNA and SPOT-RNA2 are cited from SPOT-RNA2[3].

Table S6: p-value of RNADiffFold compared with other methods on four datasets.

| Method | ArchiveII | bpRNA TS0 | bpRNA-new | PDB |
|---|---|---|---|---|
| Ufold | 6.032e-14 | 1.683e-07 | 1.214e-23 | 1.457e-01 |
| e2efold | 0.000e+00 | 0.000e+00 | 0.000e+00 | 1.137e-54 |
| MXfold2 | 4.428e-215 | 2.600e-35 | 2.297e-07 | 5.997e-01 |
| Linearfold | 0.000e+00 | 3.119e-54 | 8.125e-16 | 1.780e-02 |
| Contrafold | 0.000e+00 | 1.419e-52 | 3.239e-05 | 2.489e-01 |
| Mfold | 0.000e+00 | 1.296e-77 | 8.644e-35 | 5.310e-02 |
| RNAfold | 0.000e+00 | 2.023e-70 | 1.919e-16 | 1.785e-01 |
| RNAStructure | 0.000e+00 | 2.492e-71 | 3.418e-26 | 8.492e-02 |

Table S7: 95% confidence intervals obtained by the bootstrap percentile method for PDB dataset with various bootstrap steps.

| Bootstrap steps | TS1 | TS2 | TS3 |
|---|---|---|---|
| 20 | (0.689, 0.712) | (0.808, 0.821) | (0.674, 0.689) |
| 50 | (0.695, 0.710) | (0.805, 0.824) | (0.675, 0.690) |
| 100 | (0.691, 0.713) | (0.806, 0.823) | (0.673, 0.692) |
| 200 | (0.690, 0.717) | (0.804, 0.821) | (0.674, 0.690) |
| 500 | (0.690, 0.715) | (0.805, 0.823) | (0.673, 0.692) |
| 1000 | (0.690, 0.716) | (0.805, 0.822) | (0.672, 0.691) |
| 2000 | (0.690, 0.715) | (0.806, 0.823) | (0.673, 0.691) |
| 5000 | (0.690, 0.715) | (0.805, 0.823) | (0.672, 0.691) |

Table S8: F1, precision and recall of RNADiffFold on four test sets given num sample=1 and 10.

| Test set | num sample=1 | | | num sample=10 | | |
|---|---|---|---|---|---|---|
| | Prec | Rec | F1 | Prec | Rec | F1 |
| RNAStrAlign test | 0.952 | 0.946 | 0.948 | 0.955 | 0.949 | 0.951 |
| ArchiveII | 0.886 | 0.873 | 0.875 | 0.889 | 0.878 | 0.880 |
| bpRNA TS0 | 0.701 | 0.739 | 0.702 | 0.717 | 0.743 | 0.711 |
| bpRNA-new | 0.661 | 0.686 | 0.652 | 0.672 | 0.693 | 0.659 |

Table S9: Ablation studies on ArchiveII, bpRNA TS0, bpRNA-new and PDB datasets. v1-v5 are different condition construction processes combined with the diffusion process. v1: one-hot encoding $c_{onehot}$; v2: probability map from the Ufold score network $c_u$; v3:one-hot encoding $c_{onehot}$, output from RNA-FM $c_{emb}, c_{attn}$; v4, one-hot encoding $c_{onehot}$, probability map from the Ufold score network $c_u$, output from RNA-FM $c_{emb}$; v5, full version of RNADiffFold.

| | ArchiveII | | | bpRNA TS0 | | | bpRNA-new | | | PDB | | |
|---|---|---|---|---|---|---|---|---|---|---|---|---|
| | Prec | Rec | F1 | Prec | Rec | F1 | Prec | Rec | F1 | Prec | Rec | F1 |
| v1 | 0.904 | 0.887 | 0.892 | 0.632 | 0.654 | 0.627 | 0.579 | 0.591 | 0.561 | 0.779 | 0.510 | 0.601 |
| v2 | 0.860 | 0.827 | 0.837 | 0.667 | 0.701 | 0.656 | 0.664 | 0.669 | 0.645 | **0.891** | 0.577 | 0.684 |
| v3 | **0.909** | **0.910** | **0.906** | 0.627 | 0.724 | 0.660 | 0.562 | 0.666 | 0.598 | 0.770 | 0.606 | 0.668 |
| v4 | 0.870 | 0.841 | 0.851 | 0.675 | **0.744** | 0.691 | 0.651 | **0.725** | **0.672** | 0.696 | **0.746** | 0.713 |
| v5 | 0.889 | 0.878 | 0.880 | **0.717** | 0.743 | **0.711** | **0.672** | 0.693 | 0.659 | 0.868 | 0.653 | **0.736** |

Table S10: The hyper-parameters configuration.

| Hyper-parameter name | Value | Description |
|---|---|---|
| epoch | 400 | The preset epochs for training |
| batch size[a] | 1 | The input batch size after data bucketing |
| diffusion steps | 20 | The number of steps for diffuse and denoise |
| embedding dimension | 8 | The dimension used to represent two categories of bb relationship |
| learning rate | 0.001 | The learning rate of Adam schedular |
| Early stop patience | 5 | The patience of early stop |

[a] Thanks to the data bucketing strategy, the "1" batch size contains data that is similar in length and appropriate to memory.

Table S11: Comparison of the F1 scores of RNADiffFold, Combination_1 and Combination_2 on four datasets. For detailed experimental settings, see the Supplementary Section 1.2.

| Method | ArchiveII | bpRNA TS0 | bpRNA-new | PDB |
|---|---|---|---|---|
| RNADiffFold | 0.880 | 0.711 | 0.659 | 0.736 |
| Combination_1 | 0.490 | 0.545 | 0.585 | 0.614 |
| Combination_2 | 0.737 | 0.581 | 0.598 | 0.707 |

Table S12: Comparison of different deep learning models in terms of inference time and number of parameters.

| Method | Inference Time on RNAStralign test set (Time per seq) | Number of Parameters |
|---|---|---|
| RNADiffFold | 1-2s (GPU: A40) | 836,423 |
| Ufold (Pytorch) | 0.16s (GPU) | 8,641,377 |
| Mxfold2 (Pytorch) | 0.31s (GPU) | 47,346 |
| E2EFold (Pytorch) | 0.40s (GPU) | 718,863 |
| SPOT-RNA (Pytorch) | 77.80s (GPU) | 7,759,445 |
| CDPfold (tensorflow) | 300.107s | - |
| LinearFold (C++) | 0.43s | - |
| Eternafold (C++) | 6.42s | - |
| RNAsoft (C++) | 4.58s | - |
| Mfold (C) | 7.65s | - |
| RNAStructure (C) | 142.02s | - |
| RNAfold (C) | 0.55s | - |
| CONTRAfold (C++) | 30.58s | - |

### 3 Supplementary Figures

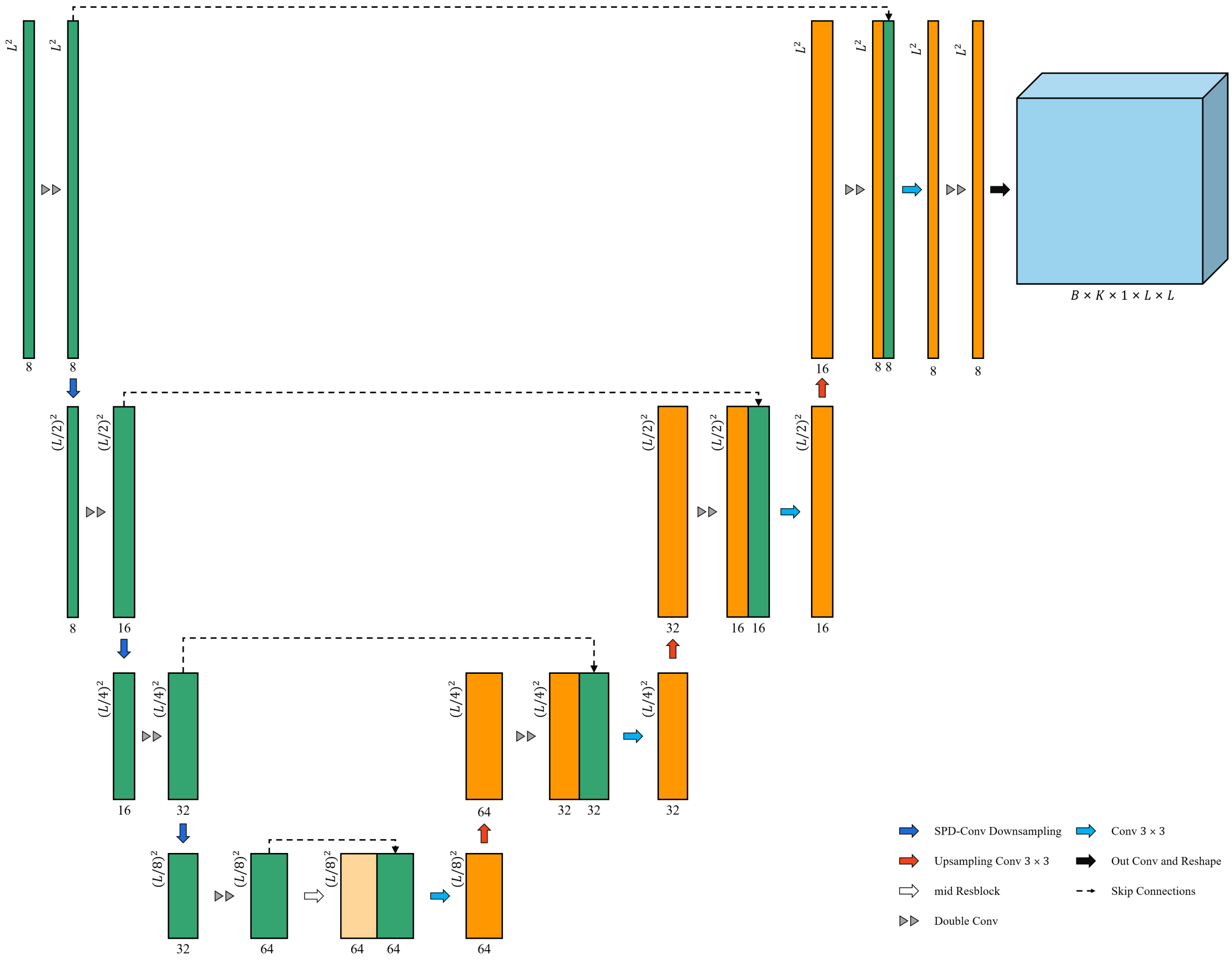


Fig. S1: The architecture of denoising Unet network

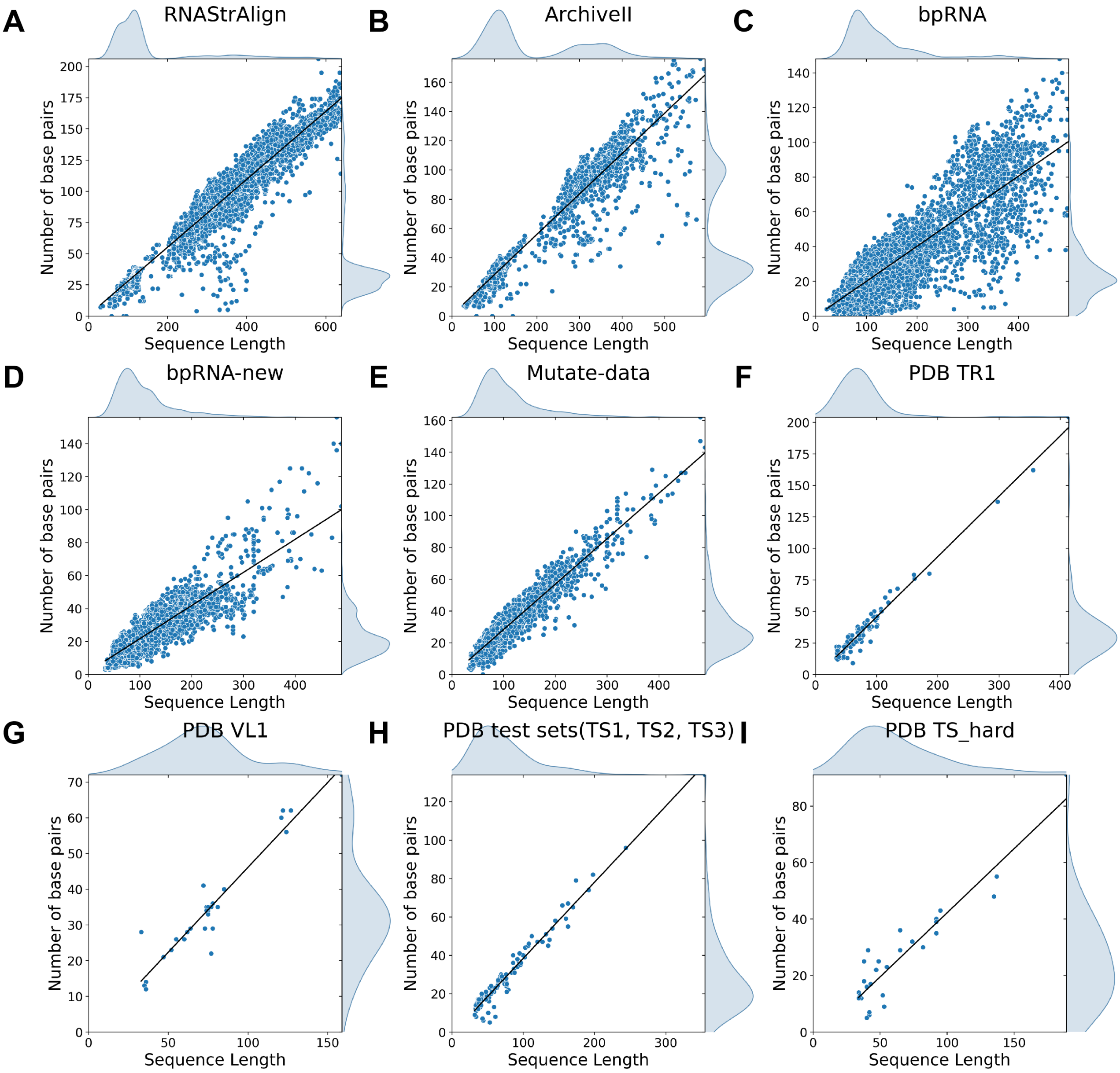


Fig. S2: The scatter plot, accompanied by marginal kernel density estimate plots, encompasses all datasets utilized, namely (A) RNAStrAlign, (B) archiveII, (C) bpRNA, (D) bpRNA-new, (E) Mutate-data, (F)PDB TR1, (G)PDB VL1, (H)PDB test sets(TS1, TS2, TS3) and (I)TS-hard. The x-axis denotes sequence length, while the y-axis represents the number of base pairs. The plot notably indicates that certain sequences exhibit a low ratio of the number of base pairs to the sequence length.

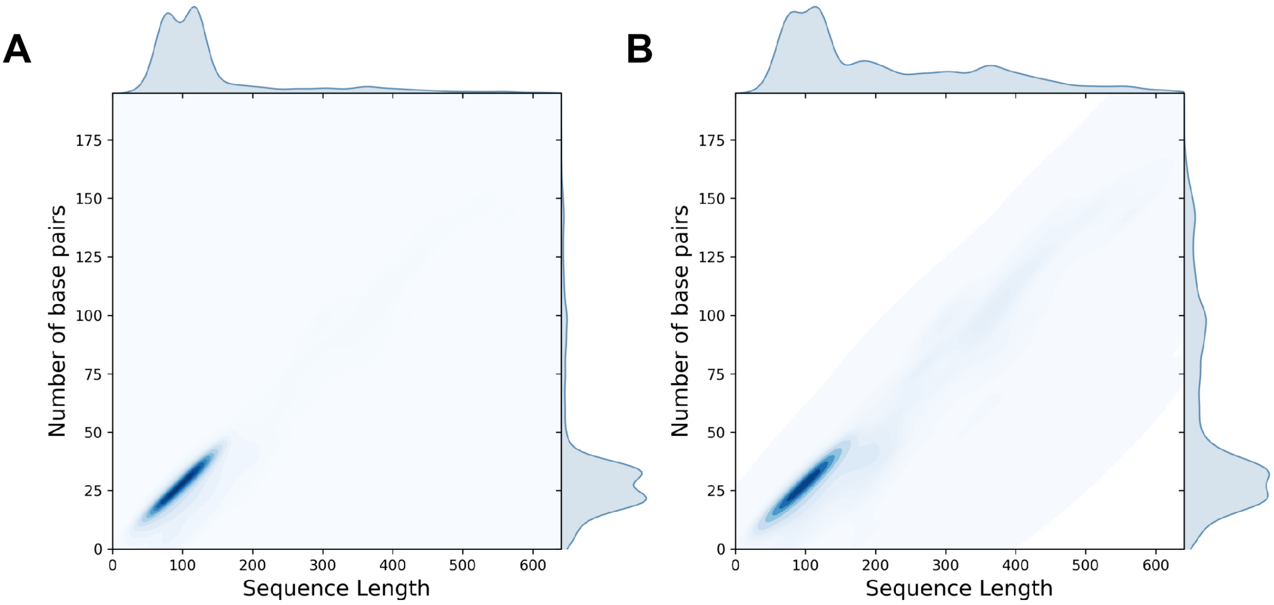


Fig. S3: The distribution of sequence length and number of base pairs before(**A**) and after(**B**) data balance on training datasets(RNAStrAlign training set, bpRNA TR0, mutate data).

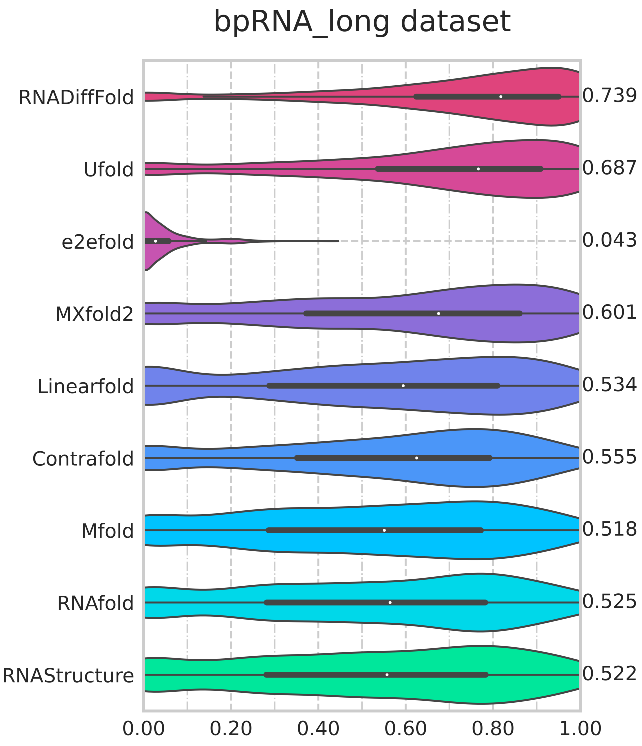


Fig. S4: long range pairing benchmark result result on TS0 dataset. Visualization of F1 value of long-range base pair prediction of RNADiffFold against 8 other methods.

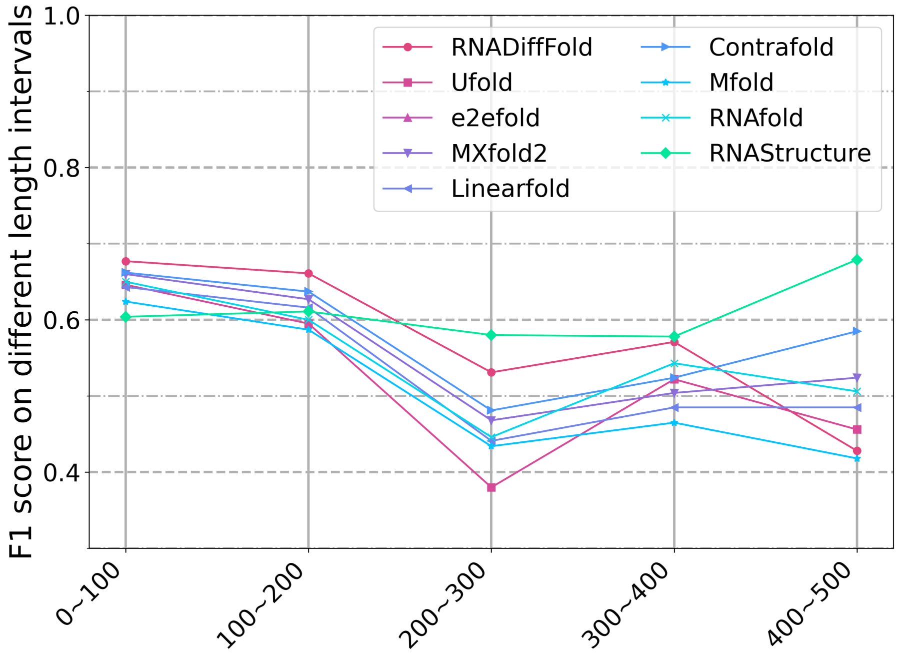


Fig. S5: Means of F1 values for different length intervals on the bpRNA-new data set. Visualization of RNADiffFold compared with other methods.

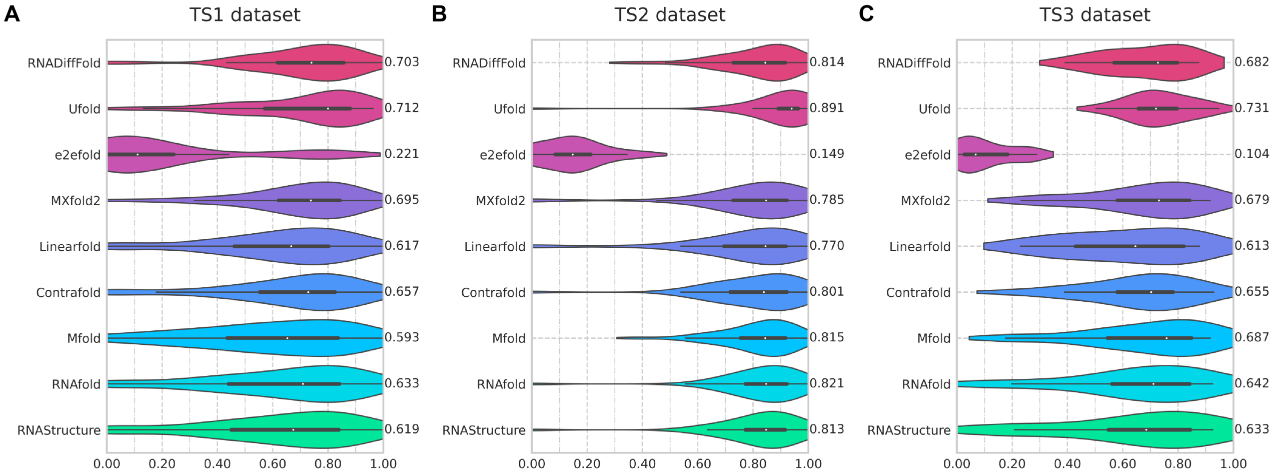


Fig. S6: Violin plot on the PDB test sets(TS1, TS2, TS3). Visualization of F1 value of RNADiffFold against 8 onther methods.

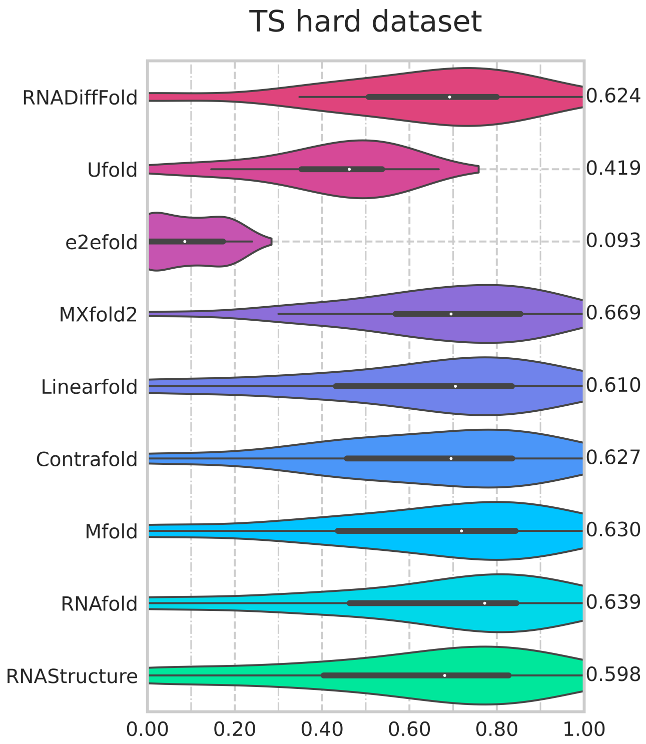


Fig. S7: Violin plot on the TS hard dataset. Visualization of F1 value of RNADiffFold against 8 onther methods.

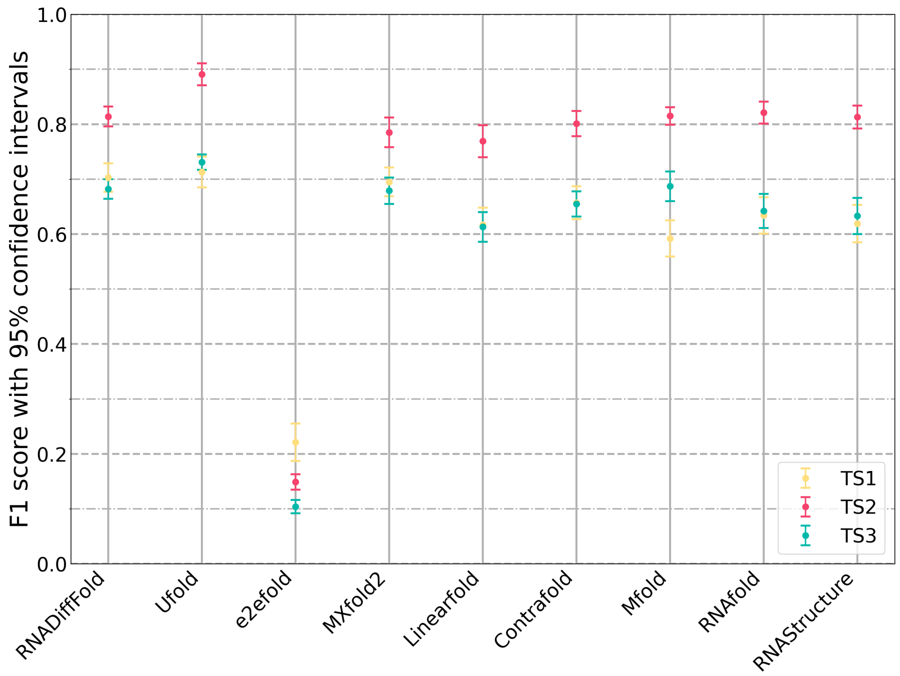


Fig. S8: 95% bootstrap percentile confidence intervals for the F1 value of all the compared methods on PDB dataset(TS1, TS2, TS3).

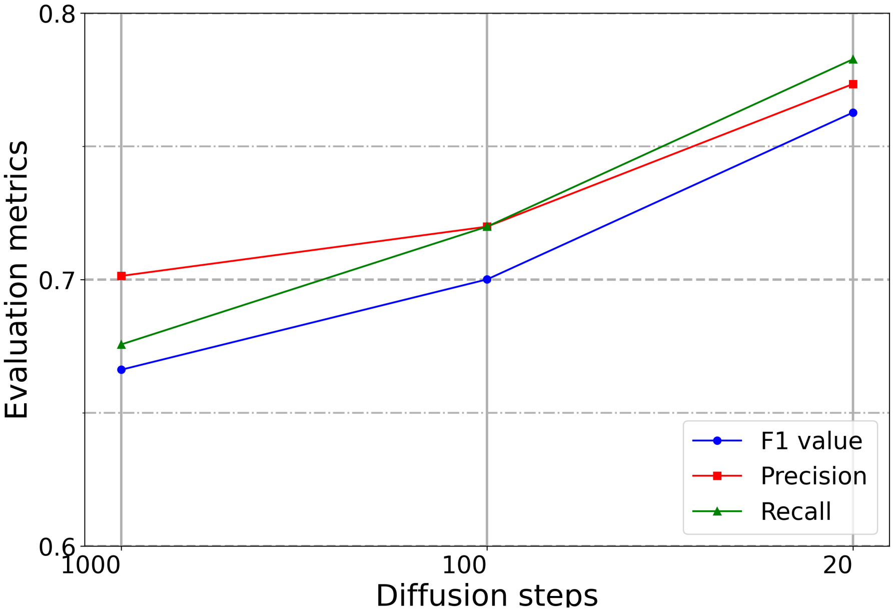


Fig. S9: Impact of the diffusion steps on bpRNA TS0(<160nt).

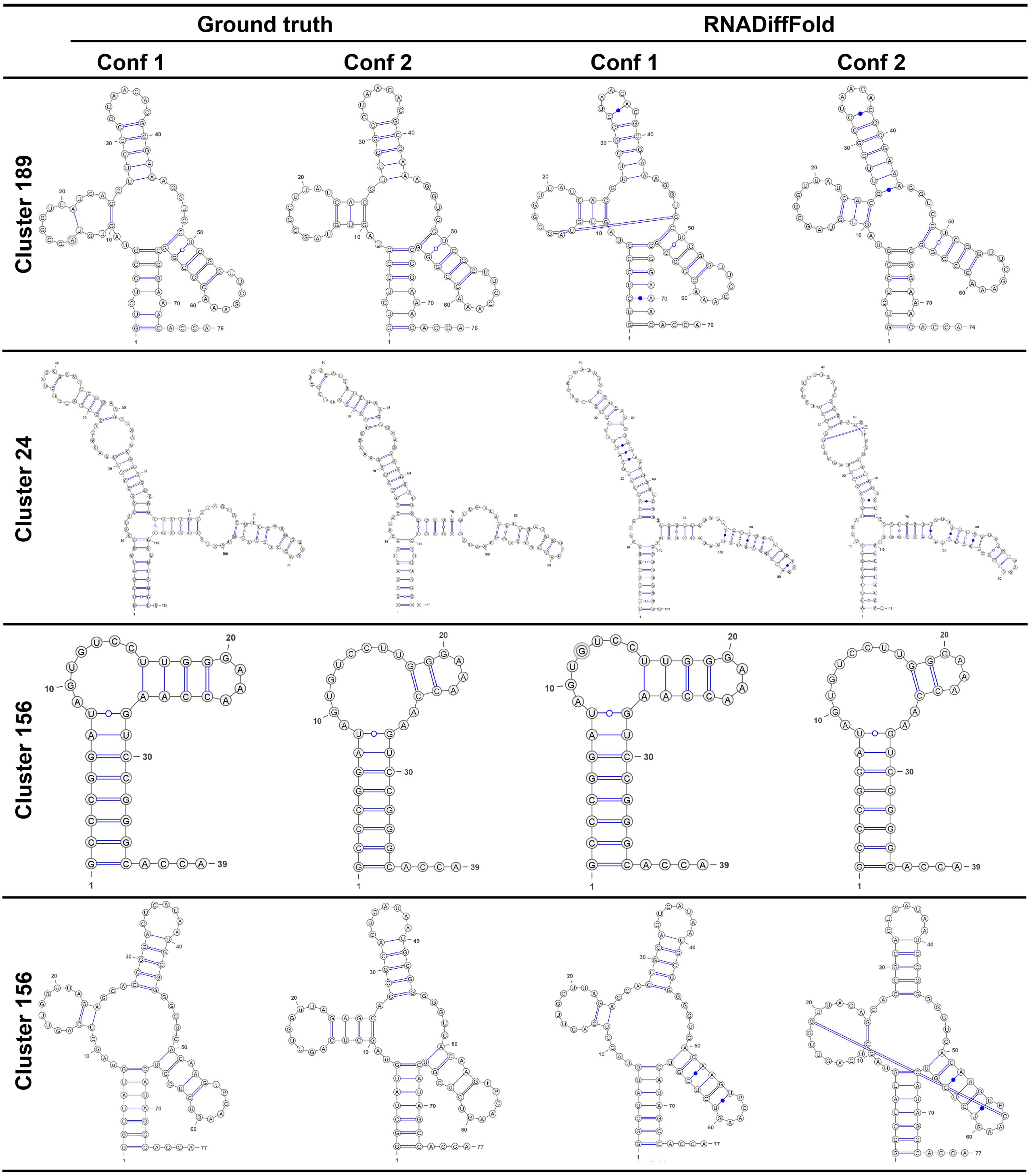


Fig. S10: Visualization of secondary structures for 4 clusters out of the 11 successfully predicted results. "**conf1**" and "**conf2**" represent two different conformations of the same sequence.

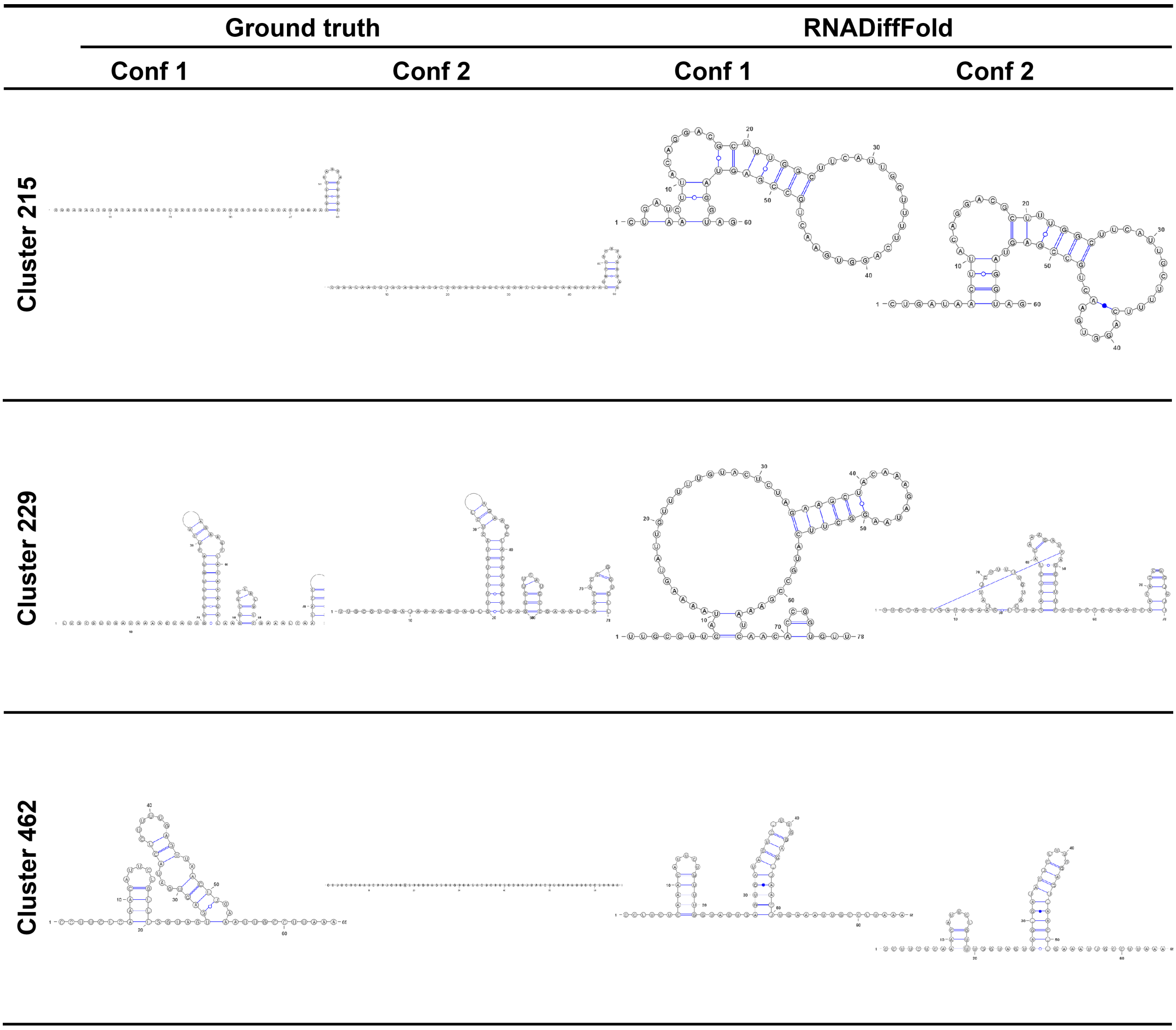


Fig. S11: Three failed cases from 14 predicted clusters. Note that the ground truth structures have low quality.

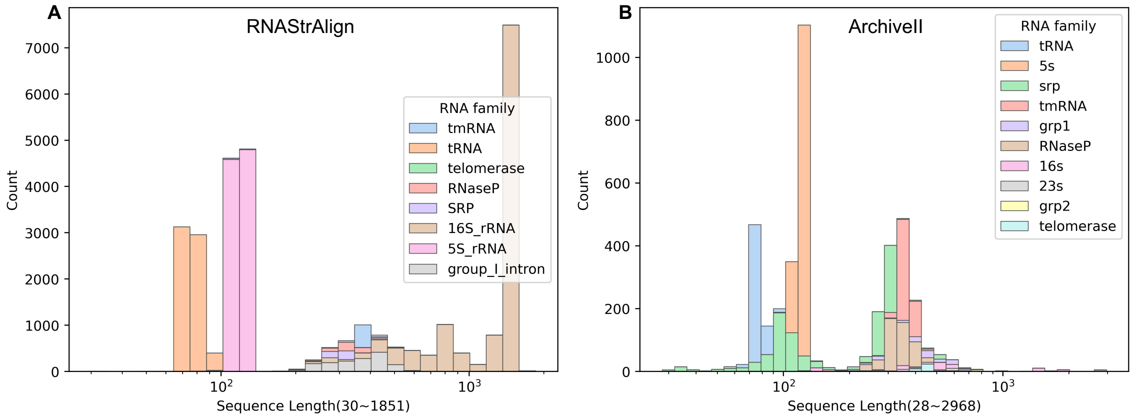


Fig. S12: Statistical analysis results of RNA sequences and family distribution on RNAStrAlign and Archive II datasets.

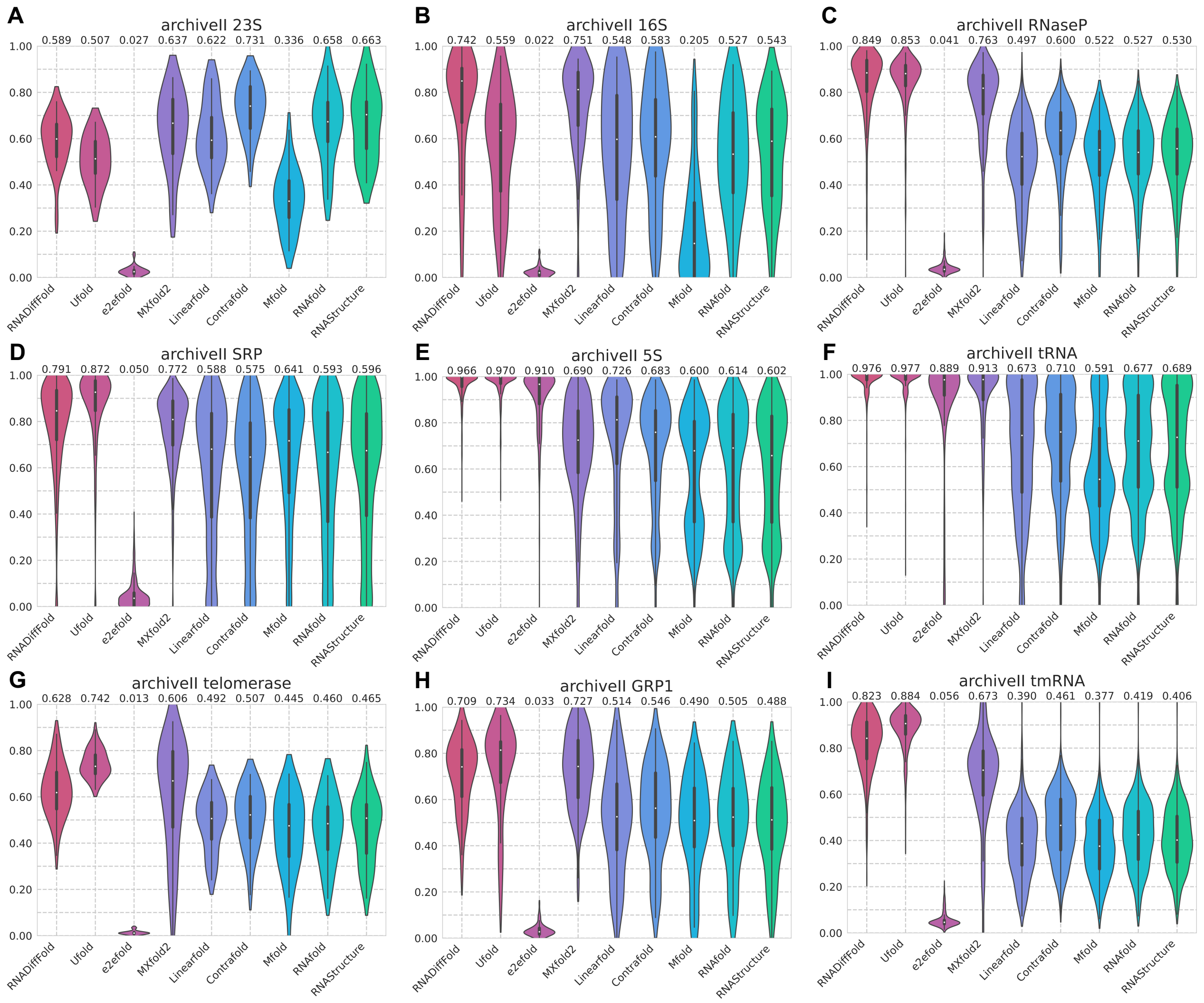


Fig. S13: This figure shows the analysis results of the predictions made by various models on the ArchiveII dataset, categorized by RNA family type. Due to the sequence lengths in the grp2 family exceeding 600 nucleotides, they were filtered out during data preprocessing, and thus the predictive results for this family were not included in the statistics.
